# Tumor microenvironment governs the prognostic landscape of immunotherapy for head and neck squamous cell carcinoma: A computational model-guided analysis

**DOI:** 10.1101/2024.09.26.615149

**Authors:** Priyan Bhattacharya, Alban Linnenbach, Andrew P. South, Ubaldo Martinez-Outschoorn, Joseph M. Curry, Jennifer M. Johnson, Larry A. Harshyne, Mỹ G. Mahoney, Adam J. Luginbuhl, Rajanikanth Vadigepalli

**Affiliations:** Department of Pathology and Genomic Medicine, Thomas Jefferson University, Philadelphia, PA19107, USA; Department of Otolaryngology, Sidney Kimmel Medical College, Thomas Jefferson University, Philadelphia, PA19107, USA; Department of Pharmacology, Physiology, and Cancer Biology, Thomas Jefferson University, Philadelphia, PA19107, USA; Department of Medical Oncology, Sidney Kimmel Medical College, Thomas Jefferson University, Philadelphia, PA19107, USA; Department of Microbiology and Immunology, Sidney Kimmel Medical College, Thomas Jefferson University, Philadelphia, PA19107, USA

**Author notes:** Corresponding author: Rajanikanth Vadigepalli.

**Keywords:** Tumor microenvironment, Head and neck squamous cell carcinoma, Computational modeling, Systems biology, Cancer resistance, Immunotherapy

## Abstract

Immune checkpoint inhibition (ICI) has emerged as a critical treatment strategy for squamous cell carcinoma of the head and neck (HNSCC) that halts the immune escape of the tumor cells. Increasing evidence suggests that the onset, progression, and lack of/no response of HNSCC to ICI are emergent properties arising from the interactions within the tumor microenvironment (TME). Deciphering how the diversity of cellular and molecular interactions leads to distinct HNSCC TME subtypes subsequently governing the ICI response remains largely unexplored. We developed a cellular-molecular model of the HNSCC TME that incorporates multiple cell types, cellular states, and transitions, and molecularly mediated paracrine interactions. An exhaustive simulation of the HNSCC TME network shows that distinct mechanistic balances within the TME give rise to the five clinically observed TME subtypes such as immune/non-fibrotic, immune/fibrotic, fibrotic only and immune/fibrotic desert. We predict that the cancer-associated fibroblast, beyond a critical proliferation rate, drastically worsens the ICI response by hampering the accessibility of the CD8+ killer T cells to the tumor cells. Our analysis reveals that while an Interleukin-2 (IL-2) + ICI combination therapy may improve response in the immune desert scenario, Osteopontin (OPN) and Leukemia Inhibition Factor (LIF) knockout with ICI yields the best response in a fibro-dominated scenario. Further, we predict Interleukin-8 (IL-8), and lactate can serve as crucial biomarkers for ICI-resistant HNSCC phenotypes. Overall, we provide an integrated quantitative framework that explains a wide range of TME-mediated resistance mechanisms for HNSCC and predicts TME subtype-specific targets that can lead to an improved ICI outcome.

## Introduction

Squamous cell carcinoma of the head and neck (HNSCC) is a collection of head and neck cancers typically initiated from the mucosal cells in the larynx, pharynx, and oral cavity (1,2). Recent clinical observations suggest that the response to immunotherapy strongly correlates with the cellular composition of the associated tumor microenvironment (TME) (3,4). These observations necessitate elucidating the non-tumor cells’ precise role in the survival and growth of tumor cells. The existing targeted therapies focusing on the intrinsic signaling factors within the tumor cells underplay the prospect of the TME functioning as a ‘networked system’ with emergent properties (5–7). On the other hand, immunotherapies such as immune checkpoint inhibitors (ICI) attempt to strengthen the immune response of the TME through the elimination of specific molecular programs (such as programmed death-1 (PD1)/ programmed death ligand-1 (PDL1) or α-CTLA4) crucial for the immune escape of tumor cells (8). Although the ICI treatments, under certain conditions, yield favorable outcomes, the majority of the HNSCC patients remain non-responsive to ICI treatment, indicating the need for the analysis of the interactions of the TME components (tumor, immune, and fibrotic) towards a comprehensive understanding of the possible mechanisms for insensitivity to immunotherapy.

Mathematical modeling of the tumor cell population can provide a quantitative understanding of the role of TME constituents and estimate the patient-specific anti-cancer dosage amount (9). Swan (1995) proposed that designing the chemo-dosage to reduce the tumor size with minimum side effects (loss of healthy tissue) amounts to solving an optimal control problem (10). Although recent studies have proposed model-free mechanisms for solving data-driven optimal control problems, they lack explainability and fail to answer pertinent questions about understanding therapy resistance (10,11).

Depending on the granularity, the existing mathematical models can be divided into at least three distinct categories— i) gene level, ii) cell population-based, and iii) cell-state transition networks. The gene-level modeling begins with the underlying gene regulatory network of the cell in consideration (tumor cells in this context) (12–14). The multi-state nature of the dynamical system constituted from modeling the gene regulatory networks

points to the existence of different cell states and their distinct responses to therapeutic strategies (15). In a stochastic setting, a non-zero transition probability (governed by mutations and intercellular interactions) may exist for each trajectory in a particular cell state’s neighborhood to gravitate toward a different cell state (12, 16-19). Modeling a gene regulatory network requires dealing with a large dimensional system with highly non-linear dynamics, making the model less intuitive and inconvenient for theoretical interventions. Extending the scope of the gene-level analysis to the ‘non-tumor’ cells is computationally taxing due to the large dimension of the system. Therefore, although the gene-level analysis inspires novel targeted therapies focused on the tumor cells, a TME-wide understanding may not be feasible with this framework.

Cell-population-based networks overcome the dimensionality issue at gene levels by modeling the interaction between different cell types and the response of tumor cells to anti-tumor therapies (20–23). Cell-level modeling primarily focuses on analyzing the response to a particular treatment regime. In chemotherapy-based modeling, the cell-level models involve the external drugs and the tumor cell population (20,24). In contrast, an immune-therapy-centric model revolves around the killer T-cells and the tumor cells (22, 23). The cell population-based models do not capture the inherent complexity in reliable detail. For instance, as discussed above, each cell type can have multiple distinct cell states with functionally opposite effects on the growth of the tumor cells. Cell-type population modeling cannot capture this inherent complexity, thereby failing to explain some peculiar therapeutic outcomes.

The development of high-throughput, single-cell transcriptomic analysis has made it possible to identify discrete cellular states. Further, the existing biological knowledge and transition path theory-based algorithms are also used to create conversion links between the cellular states, thereby paving the way for a cell-state-based network representation (16). The cell state models begin with different cellular states of a given cell type. The cellular states for each cell type can transition among themselves in the presence of specific biological promoters (25). For instance, in the epithelial-to-mesenchymal transition (EMT) program, the epithelial cell states convert to mesenchymal cells in the presence of several pro-inflammatory cytokines (26). Further, two distinct cell states from different cell types can affect the proliferation of each other via either paracrine interactions or cytokine secretions (27–29). The possible cell states for a given cell type are evaluated from the single-cell RNA sequencing data literature. Based on the expression of marker genes, tumor cells can be divided into multiple distinct expression states depending on the type of cancer. For instance, in HNSCC, Puram et al. (2017) have identified seven different HNSCC tumor cell states, namely cycling (G2/M, G1/s), partial EMT, hypoxia, epithelial differentiation (1,2), and stressed from the single RNA sequence data of 6000 single cells collected from 18 patients (30). A similar pattern of cell state multiplicity has also been discovered from melanoma patients’ single-cell RNA sequence analysis (4). Like tumor cells, single-cell datasets also indicate multiple cell states for non-tumor cells such as T cells, fibroblasts, and macrophages.

The cell-state transition models, by design, can potentially circumvent the explainability issues encountered in cell population-based models. Further, since the number of cell states represents the cardinality of the attractor sets in the underlying regulatory dynamics, the model’s dimensionality is reduced from the gene-level models (15). Although several scholarly interventions have attempted to propose cell state models in the context of cancer, extension of the same to the entire TME for cancer (particularly HNSCC) remains an open area of study.

Considering the limitations and possibilities of different modeling paradigms, this work operates at the cell state level, wherein we reconstruct an essential interaction atlas for HNSCC based on an extensive literature survey. In the state space form, we represent the population and concentration of the cellular states and molecular species. Via an exhaustive scanning across the model parameter space, we identify five distinct stationary TME compositions that resemble the well-known, clinically verified TME subtypes, such as immune/fibrotic, immune/non-fibrotic, non-immune/fibrotic, and desert. Further, our investigation shows the immune/fibrotic composition can further be classified into two subcategories: fibro-dominated (fibro-rich) and immune-dominated (immune-rich). We observe that the quantitative balance between cancer-associated fibroblasts (CAF)-tumor interactions and the cytotoxicity of the killer T cell governs the post-ICI outcome in fibro and immune-dominated scenarios. While the immune desert TME, in most of the scenarios, remains insensitive to the ICI therapy, the immune-dominated HNSCC TME is characterized by a significant reduction of tumor cells and CAF compared to its pre-ICI population. The fibro-dominated TME, due to CAF-induced remodeling of the extracellular matrix, alters the accessibility of the killer T cells to the tumor cells. Our models suggest that a one-time IL-2-based intervention, beyond a critical T cell proliferation rate, can overcome the ICI insensitivity posed by an immune desert TME. We identify cytokines such as OPN and LIF as important targets to improve the ICI response of a fibro-dominated scenario. Finally, IL-8 and lactate show distinct signatures across different HNSCC TME subtypes identified by the model, indicating the potential to be considered essential biomarkers for therapy resistance.

## Results

We reconstruct the overall TME network structure for HNSCC from an extensive survey of the existing literature (30-90). As shown in Figure 1, from the perspective of immunotherapy (specifically anti-PD1 therapy), the tumor cells are divided into three different cellular states, namely, 1) stem, 2) without Programmed death ligand-1 (PDL1) expression (PDL1−), and 3) with PDL1 (PDL1+). The T cell modules contain three different cell types with four distinct cellular states obtained from the single cell RNA sequence (scRNA-seq) analysis of the HNSCC TME by Puram *et al. (2017)*. The CD8+ cytotoxic killer T cells, CD4+ helper T cells, CD4+ CD25+ regulator T cells, and CxCL13+ exhausted T cells. Fibroblasts are found in two different cellular states, namely, the wild-type fibroblasts that partake in either a tumor-suppressing role or remain quiescent. However, the wild-type fibroblasts, at a later stage of tumor development, may transition into an invasive phenotype that promotes the growth of tumor cells and alters the T to tumor cell accessibility. Similarly, the present work considers two different macrophage states based on their polarities. Finally, the proposed network incorporates eight different molecular species (chemokines, cytokines, and metabolites), namely-Interleukins (IL) IL-2, IL-8, IL-10, Interferon Gamma (IFNG), Osteopontin (OPN), Interferon regulatory factor 8 (IRF8), Leukemia inhibitory factor (LIF), and Lactate (Lac).

**Fig. 1.**
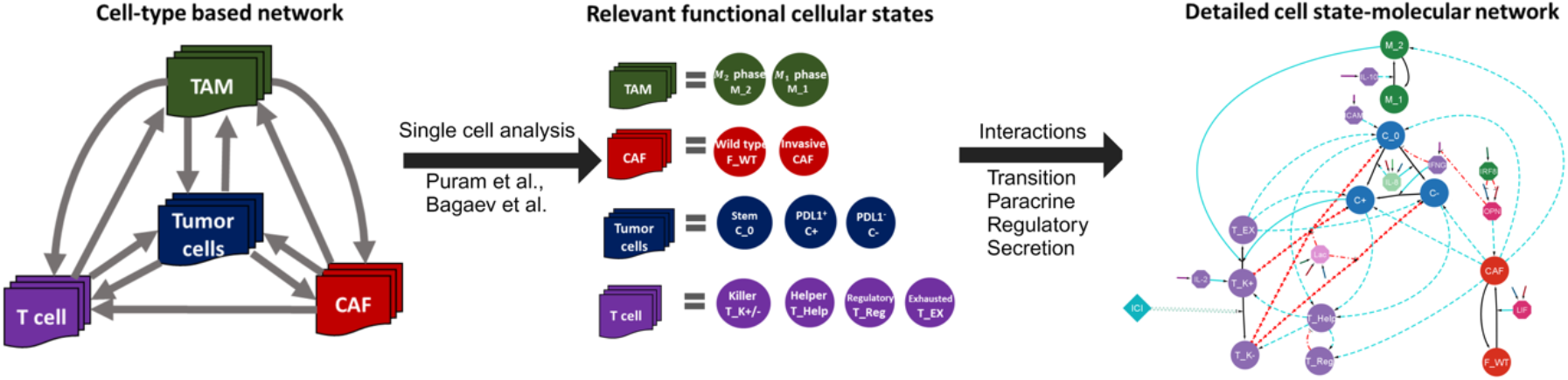
Construction of cell-state specific tumor-microenvironment network for HNSCC: We begin with the relevant cell types in a HNSCC TME obtained from the literature. The existing cell-type-specific models do not capture the interactions reliably, for mutually opposite effects may exist between two cell types (*e*.*g*., T cells and tumor cells). Subsequently, the literature on single-cell studies of HNSCC TME reveals the functionally distinct cellular states for different cell types (Puram *et al*. (2019), Bagaev *et al*. (2021)). Similarly, we searched for the possible interactions between different cell states of interest to establish a cell-state interaction network. Further, we considered relevant molecular species from the HNSCC literature that affect the population of individual cell states.

With the modeling rules and assumptions declared in the methodology section, the proposed model develops into a dynamic system with twenty-four states and ninety-one parameters. The state variables denote the population (concentration) of the different cell states (molecular species). The assumption of spatial homogeneity translates to a system of ordinary differential equations (ODE).

### Classification of TME compositions

To find out the possible TME compositions, we simulated the HNSCC TME system in Figure 1, with ten thousand different parameter sets pertaining to the growth, proliferation, conversion, and death fluxes for CAF, Killer T, tumor cell population. We have also repeated the exercise in the scenario of competitive resources for five different maximum resource capacities. Interestingly, we observed that resource competition provides a better overall outcome than fixed resource rates (Fig. S2). However, in both scenarios (fixed and competitive resources), we observe five distinct cohorts in the CAF-tumor-killer T cell landscape. The non-desert TME compositions can have two different varieties, namely, fibro-dominated, and immune-dominated. A fibro-dominated TME (cohort 1 of Fig. 2(b) and 2(c)) is identified with an elevated level of CAF population and tumor cell population compared to the killer T cells.

**Fig. 2.**
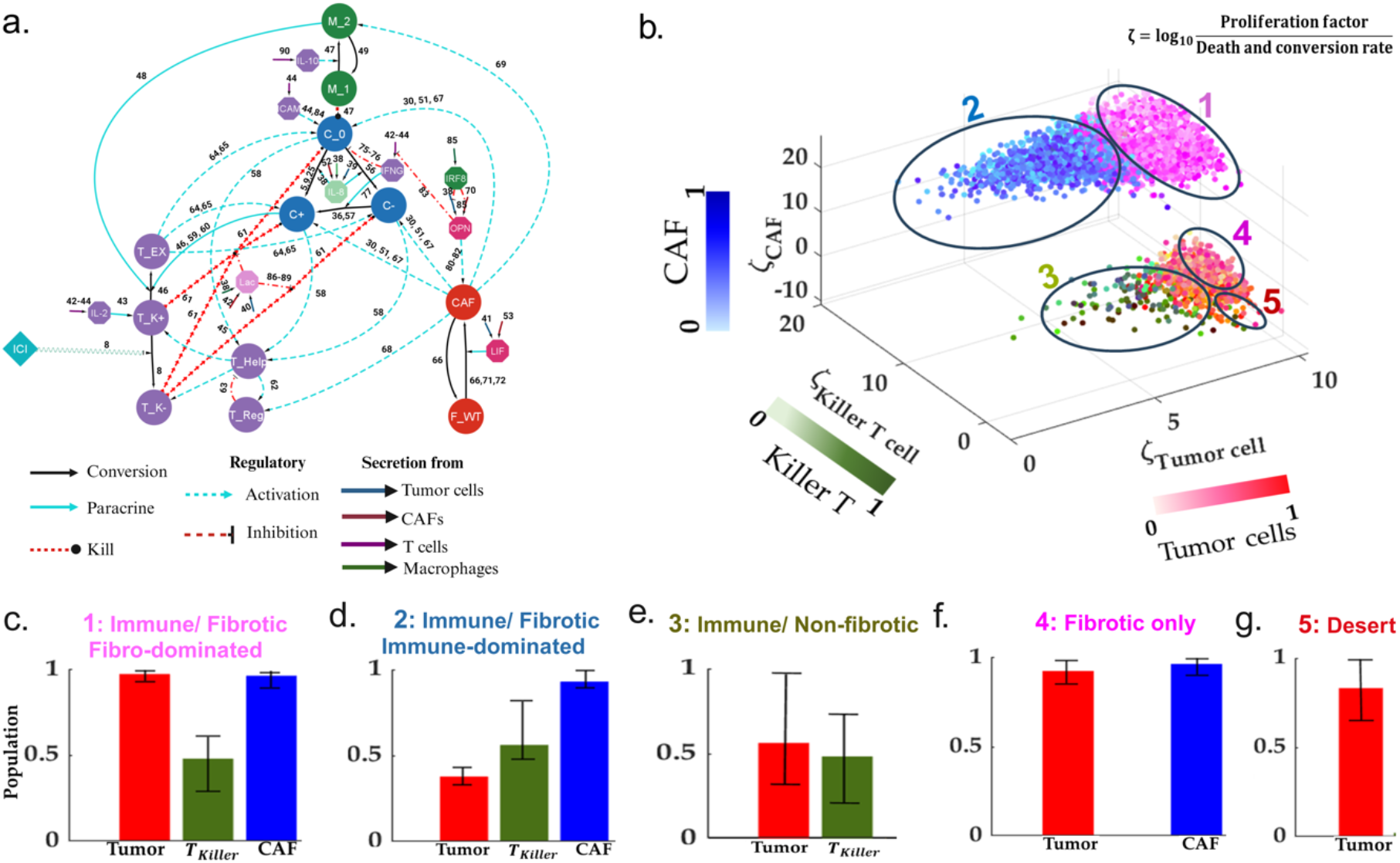
Explaining the phenotypic heterogeneity of the HNSCC TME. **(a)** The nodes are either the cell states or the molecular species, whereas the edges represent diverse forms of interactions. The acronyms C_0, C_PDL1+, and C_PDL1- refer to stem, PDL1+ (programed death ligand1), and PDL1- tumor cells, respectively. T_K+, T_K-, T_Help, T_Reg, and T_Ex stands for PD1+ (programmed death 1), PD1- killer T cells, Helper T cells, Regulatory T cells, and Exhausted T cells, respectively. M_1 and M_2 refer to macrophages of M1 and M2 phase, respectively. Further, F_WT and CAF correspond to wild-type and invasive cancer-associated fibroblasts, respectively. The acronyms IL-2, IL- 8, IL-10 LIF, IFNG, IRF8, OPN, ICAM1, and Lac denote Interleukin-2, Interleukin-8, Interleukin-10, Leukemia Inhibitory Factor, Interferon Gamma, Interferon Regulatory Factor-8, Osteopontin, Intercellular Adhesion Molecule-1, and Lactate, respectively. **(b)** We simulated the dynamics for 10,000 parameter combinations based on the reconstructed network system to obtain the possible steady-state clusters. We found five distinct clusters: desert, immune desert, fibro desert, immune-dominated, and fibro- dominated. **(c-g)** Bar charts for the median population of tumor cells, killer T cells, and CAF population across different cohorts. The lower and upper intervals indicate the 25% and 75% quartiles, respectively.

Meanwhile, the immune-dominated TME lacks the population of PDL1 tumor cells and contains a strong presence of cytotoxic T cells (cohort 2 of Fig. 2(b) and 2(d)). Interestingly, the immune and fibro-dominated TME contains similar levels of immune and CAF cells. Therefore, it is not the abundance of CAF and immune cells but the balance between the tumor-promoting role of CAF and the cytotoxicity of killer T cells that determines the tumor cell population, which serves as the crucial distinguishing factor between the two TME phenotypes. Further, the ratio between the PDL1-to PDL1+ tumor cells can be a potential biomarker for distinguishing two distinct TME phenotypes.

On the other hand, the desert phenotypes are identified as having zero or very few immune systems or (and) CAF population. Owing to the absence of the CAF population, the immune/non-fibrotic TME composition (cohort 3 of Fig. 2(b) and 2(d)) is characterized by a lower tumor cell count compared to the immune desert scenario (cohort 4 of Fig. 2(b) and 2(e)). We observe that although the immune-dominated, immune/non-fibrotic, and desert arrangements can be characterized with a lower tumor cell population compared to the fibro-dominated and immune-desert phenotypes (cohort 5 of Fig. 2(b) and 2(f)), the HNSCC TME is never completely depleted of the tumor cells in the pre-ICI setting primarily due to the resource availability and immune escape and exhaustion mechanisms specific to each of the identified stationary TME compositions.

### Diversity within the immune desert

We observe that the immune desert (desert) contributes as the primary source of high PDL1-tumor cells. Interestingly, even for a significant fraction of the fibro-dominated scenario, the final killer T cell population settles to extremely low levels despite a high proliferation-to-death balance. Further, several scenarios pertaining to the immune desert setting reveal a moderate to high exhausted and regulatory T cell population. Therefore, we simulated the TME system for varying proliferation and exhaustion rates for different T cells in the immune module to understand the diverse mechanisms of immune depletion as well as the possible compositional variants within the immune desert. Our analysis reveals that apart from a critically low proliferation rate, a strong T cell exhaustion program may lead to an immune desert phenotype (Fig 3(a-b)). Interestingly, a specific scenario may mimic the immune desert phenotype wherein the immune system is composed of many regulatory T cells with almost no cytotoxic or helper T cells. We observe that a higher paracrine interaction rate between CAFs and the regulatory T cells can trigger the immune-cold scenario despite a low proliferation-to-death ratio for the regulatory T cells (Fig 3c).

**Fig. 3.**
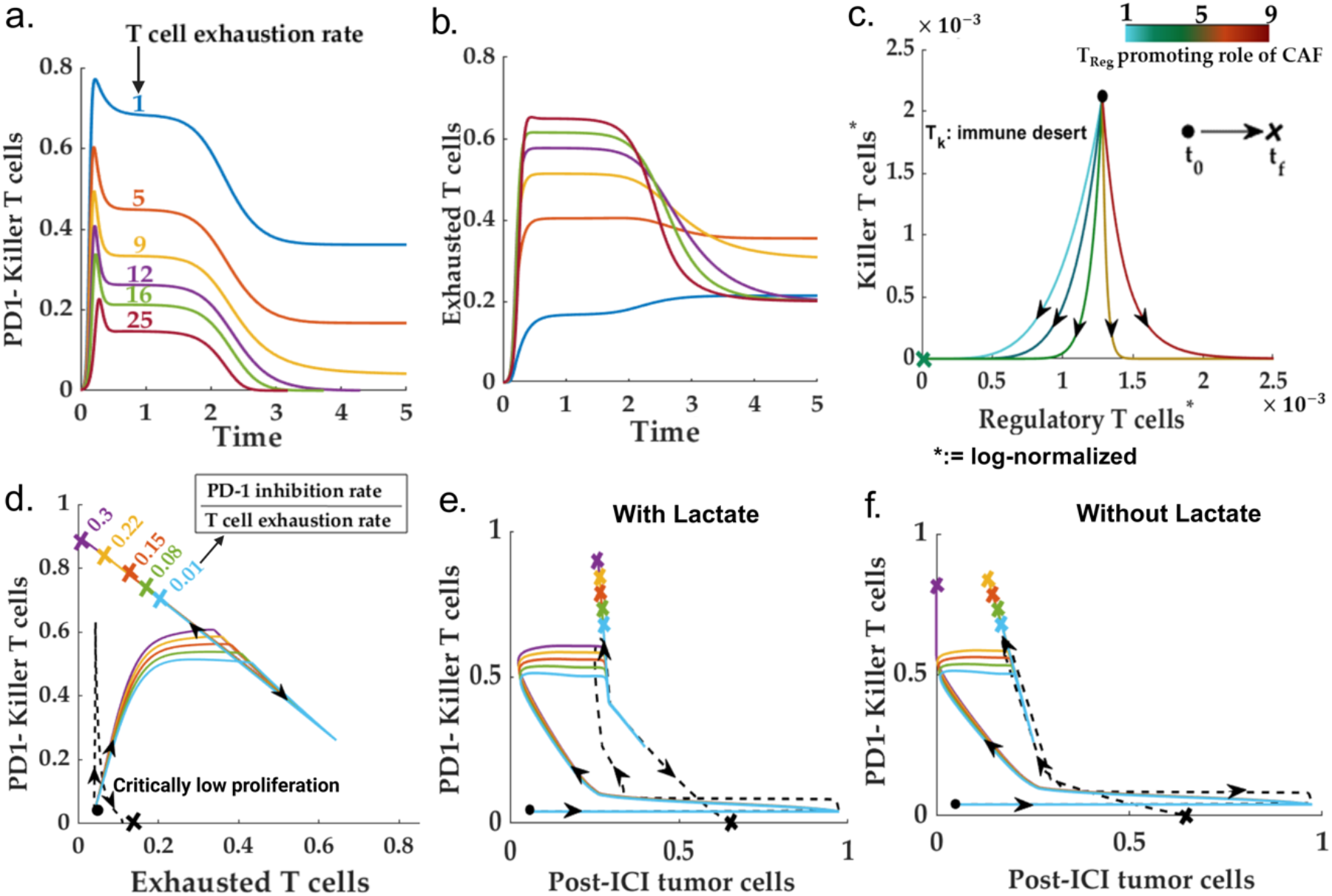
The immune-desert scenario. **(a-b)** Time profiles for killer and exhausted T cells for different T cell exhaustion rates. A high T-cell exhaustion program that can drive an immune system with a high proliferation rate to the immune-desert phenotype. **(c)** Phase space for killer T cells versus the regulatory T cells for different regulatory T cell promoting roles of CAF. Demonstrates the possibility of an immune-cold scenario with abundant regulatory T cells due to high CAF-Treg interaction. **(d)-(e)** Post-ICI trajectories of killer T cells, tumor cells, and exhausted T cells for different balances between the anti-PD1 binding rate and T cell exhaustion rate. Unlike other low-proliferation induced immune- desert phenotypes (represented in dotted lines), the ICI intervention can improve the exhaustion-driven immune desert scenario. **(f)** Lactate reduction can further reduce the total post-ICI tumor cell count.

### Effect of ICI: specific to TME subtypes?

As discussed, none of the identified stationary TME compositions leads to a complete depletion of tumor cells. The PDL1+ tumor cell population remains high in most of the scenarios—this necessitates an ICI-based intervention (anti-PD1) that eliminates the PD1 program in the killer T cells, enabling it to recognize PDL1+ tumor cells, thereby stalling the exhaustion program of the killer T cells. We studied the role of anti-PD1 dosage for all the different parametric conditions mapping to the desert (cold) and dominated TME phenotypes.

#### Are all immune deserts non-responsive to ICI?

As discussed previously, multiple mechanisms exist behind the emergence of an immune desert HNSCC TME. To investigate whether the ICI response to the immune desert scenario is dependent on the specific desert mechanism, we simulated the proposed system with a non-zero anti-PD1 level for different variants of parametric conditions reflective of all the possible cohorts within the immune-dessert phenotype in the pre-ICI scenario. To resemble a realistic scenario, the killer T cell population and IL-2 concentration are kept non-zero in the initial condition for the ICI simulation. We observed that for high binding rate of the anti-PD1 species, the exhaustion-driven immune-desert can be overcome by reducing the effective exhaustion rate whereas, a proliferation rate of the killer T cell below a critical level remains completely insensitive to ICI therapy (Figure 3(d-e)). Expectedly, like the exhausted T cells, the total tumor cell population reduces with the increase in the efficiency of the anti-PD1 therapy. However, the post-ICI in all the scenarios, leaves with a non-zero, remanent tumor cells. This is due to the low levels of effective cytotoxicity of the killer T cells. We observe that lowering residual lactate levels can improve the performance of the ICI therapy (Figure 3(f)). Moreover, our findings suggest that the responsiveness of immune deserts to ICI therapy is not uniform and depends on the specific desert mechanism. This understanding can potentially guide the development of more personalized and effective ICI therapies.

#### Immune/non-fibrotic presents a favorable scenario for ICI

The immune/non-fibrotic scenario is identified by the parametric conditions that drive the CAF population to zero in the pre-ICI setting. The pre-ICI, zero CAF population rules out the possibility of the formation of immune-inaccessible pockets inside the TME. Further, due to the absence of the most dominating interaction from CAF to TAM, a immune/non-fibrotic scenario may also be identified with a low M2 macrophage population (91). Therefore, the primary TME constituents impacting tumor cell growth are the immune cell states. For this scenario, we simulated the proposed model for varying rates of oncogenic roles of exhausted tumor cells and killer T cell cytotoxicity. A high ratio of the interaction rate between the exhausted T cell and tumor cells to the cytotoxicity of the killer T cells enhances the total tumor cell population in the pre-ICI setting. Further, this increase in the total tumor cell population is fueled by the increase in the population of PDL1+ tumor cell population (Figures 4(a)-4(b)).

**Fig. 4.**
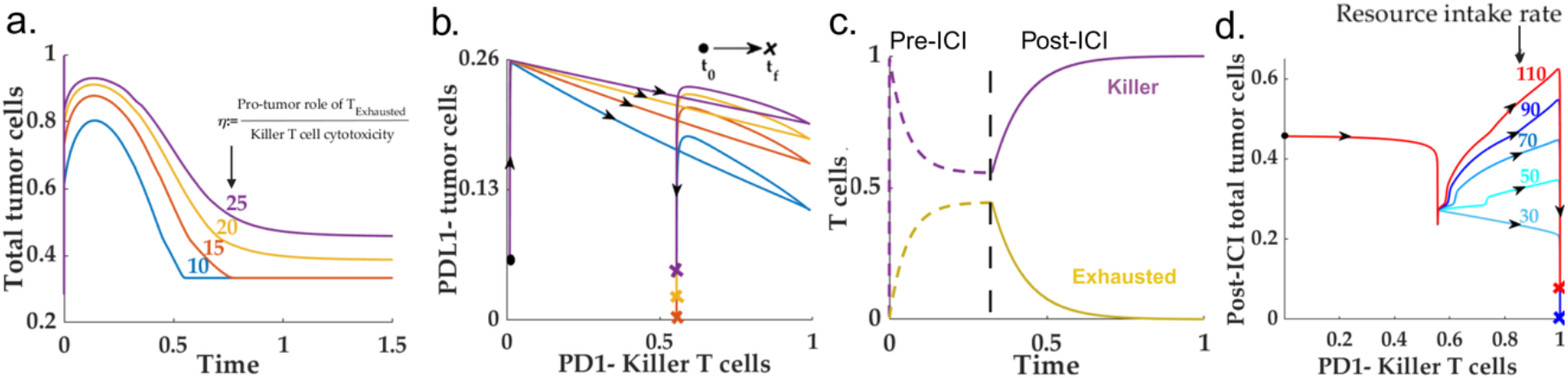
Immune/non-fibrotic, favorable case. **(a-b)** Shows despite the absence of CAF, the pre-ICI PDL1- tumor cells critically depend on the balance between the oncogenic (via Texh) and cytotoxic activities of T cells. **(c)** The intervention of ICI drives toward an aggressive cytotoxic T-cell-driven immune response. **(d)** Indicates the pivotal role of residual fixed resource supply rate (nutrients, blood flow, oxygen supply) in governing the possibility of recurrence despite a favorable prognosis.

The ICI intervention predictably yields an improved prognosis with a significant shrinkage in the tumor cell population. With a prominent level of cytotoxicity, the anti-PD1 intervention can lead to a zero exhausted T cell population (Fig 4(c)). Application of anti-PD1 favors the balance towards the tumor suppression, thereby steering the resultant total tumor cell population towards zero. Although anti-PD1 yields a favorable outcome for the immune/non-fibrotic scenario, our simulation indicates that the post-ICI scenario does not necessarily lead to a zero-tumor cell population for high resource availability (Figure 4(d)) --- This can lead to the recurrence of the disease in a post-ICI scenario. Note that due to the absence of CAFs, the growth of tumor cells occurs at a much longer timescale when compared to a fibro-dominated situation.

#### ICI reduces the CAF population in immune-dominated HNSCC TME

As discussed in the previous sections, the immune-dominated TME subtype can be identified with an abundant killer T cell population and a low PDL1-to PDL1+ tumor cell population. Therefore, the primary potential factors governing the growth, immune escape (in pre-ICI condition), and the efficacy of the ICI treatment are the killer T cells, exhausted T cells, and resident CAF population. To determine the pro-tumor components’ dependence in the HNSCC TME on the resident killer T cell population, we simulated our model with an increasing killer T cell proliferation rate. Interestingly, we found out that beyond a critical proliferation rate, the final population of killer T cells does not vary significantly with respect to the proliferation rate (Fig.5(a)). This is due to the insensitivity of the CAF population to the proliferation rate of the killer T cells (Fig. S3). Further, the CAF population remains unchanged across the killer T cell proliferation rate—This may explain the CAF abundance in an immune-dominated TME. On the other hand, for high levels of cytotoxicity, the balance between the tumor cells vis-a-vis the PDL1 signature remains heavily skewed towards the PDL1+ tumor cells (Fig.5(b)). Additionally, in resource competition environment, high levels of resource availability can be identified with high helper T cell counts owing to the abundance of PDL1-tumor cells. Our simulation suggests existence of a threshold maximum resource holding beyond which the helper T cell population becomes independent of the balance between the opposing roles of CAF (Γ) vis-a-vis the helper and killer T cell population (Fig. S4).

To determine how the exhaustion of the T cell intervenes in the immune response in a pre-ICI setting, we simulated the immune-dominated TME system with different cytotoxic levels and repeated the exercise for different exhaustion rates. Interestingly, the PDL1- - tumor cell population critically depends on the exhaustion rate of the killer T cells, but in the presence of highly cytotoxic killer T cells, the pre-ICI tumor cell population does not possess any significant dependence on the proliferation rate of the killer T cells (Figure 5(c)).

**Fig. 5.**
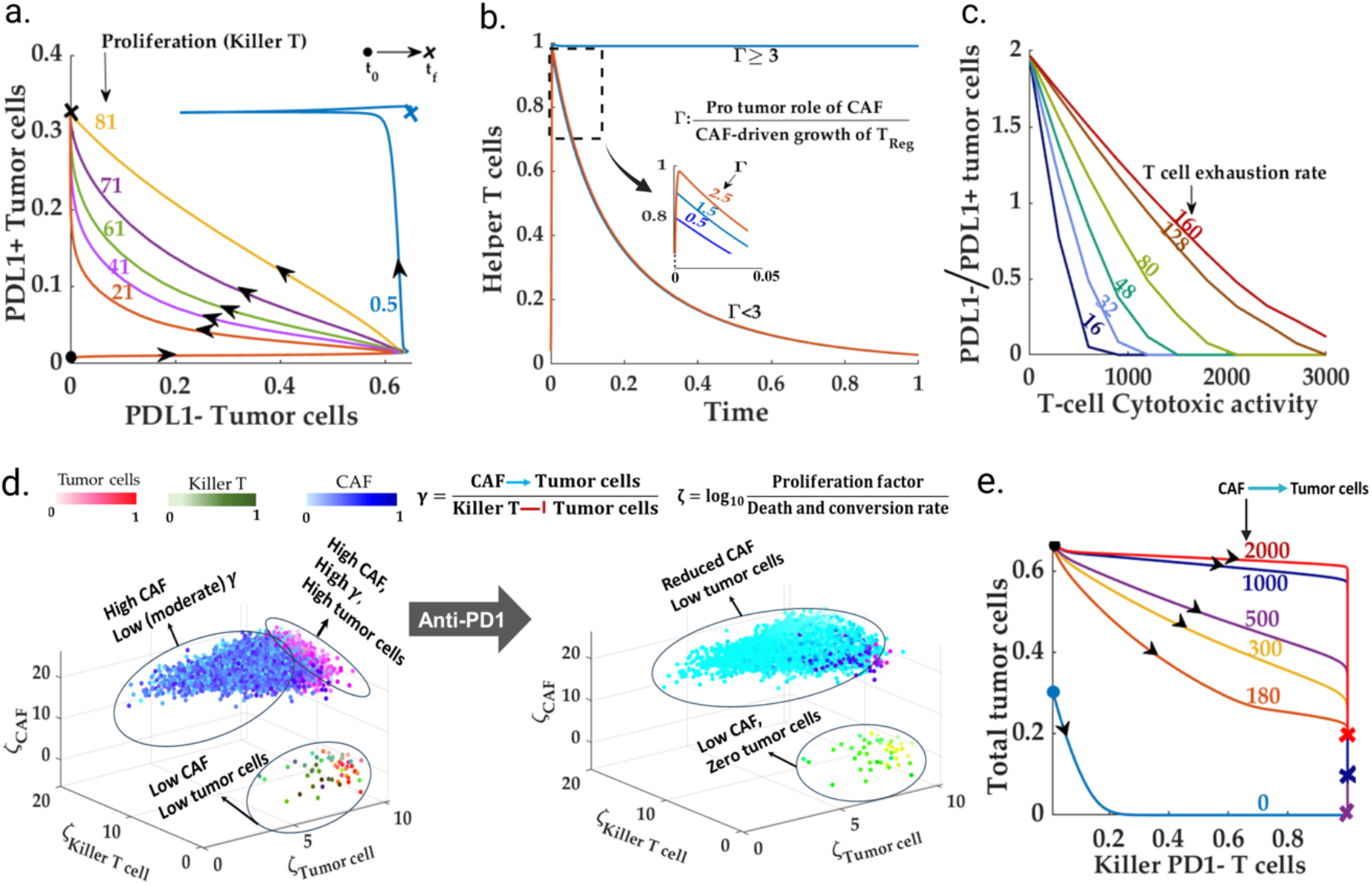
Immune-dominated-- High immune accessibility, moderate CAF: For a given CAF-tumor interaction rate, lower T-cell cytotoxicity leads to higher PDL1^-^ to PDL1^+^ tumor cell population. **(a)** For high levels of cytotoxicity, the tumor cell population remains insensitive to the proliferation rate 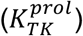 of the killer T cells in the immune dominated scenario. **(b)** CAF plays a dual role in governing the helper T cell population. CAF reduces the helper T cells via the regulatory T cells. However, a high tumor-promoting role of CAF increases the resident PDL1- -tumor cells that, in turn, increases the helper T cells via antigen sensing mechanism. **(c**) A high exhaustion rate can potentially increase the share of the PDL1- tumor cell population owing to the abundance of the exhausted tumor cells and their tumor-promoting effects. (**d)** The post-ICI scenario significantly reduces the tumor cells and CAF. The application of anti-PD1 reduces the resident CAF population and drastically reduces the tumor cell population. (**e)** The post-ICI tumor cell population is proportional to the tumor-promoting role (via paracrine interaction) of the remaining CAF population.

To examine the efficacy of an ICI-based treatment for immune-dominated subtypes, we selected 10,000 different parameter sets corresponding to the immune-dominated subtype. We simulated both the pre-ICI and post-ICI scenarios. We observed that for high immune accessibility, the immune-dominated TME subtypes significantly reduce the total tumor cell population along with the CAF population (Fig 5(d)). Although the post-ICI outcomes for immune-dominated phenotypes can be identified with a significant reduction in the CAF or, equivalently, an increase in the wild-type fibroblasts, the chance of recurrence persists due to non-zero CAF-tumor cell interaction flux owing to non-zero OPN and CAF levels (Fig. 5(e), Fig. S5). Therefore, targeting the CAF population can also be a potential therapeutic strategy in an immune-dominated TME.

#### Fibro-dominated: Effective immune desert?

Unlike the immune-dominated subtypes, the fibro-dominated TME is identified by exceedingly high CAF-tumor cell paracrine rates, low immune accessibility, and an abundant CAF population. As illustrated in Fig. 6(A), an HNSCC TME with low immune accessibility index can be conceived as dividing the TME locally into two distinct regions vis-a-vis the tumor cells—1) immune-accessible tumor cells and 2) CAF-protected tumor cells (Fig. 6(a)).

**Fig. 6.**
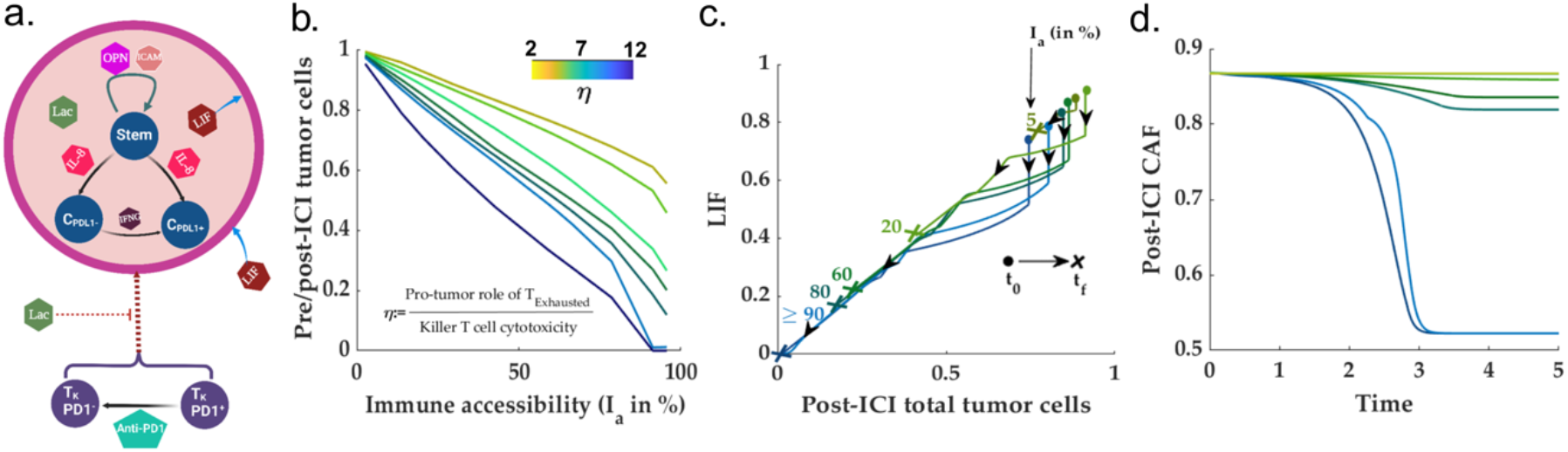
Fibro dominated, a modified immune desert. **(a)** depicts the conceptual framework of the proposed hypothesis on the CAF-pockets protected from the immune response. **(b)** demonstrate the fact that the immune accessibility index governs the post-ICI proportion of immune-accessible and immune-inaccessible tumor cells. **(c)** Phase-space for LIF and total tumor cell population. The change in LIF levels due to the ICI therapy reduces with respect to decreasing accessibility. **(d)** Contrary to the immune-dominated TME, the post-ICI CAF population remains almost constant at low immune accessibility due to a lack of change in the post-ICI LIF levels compared to its pre-ICI counterpart.

To find out the dependence of the overall ICI outcome on immune accessibility, we simulated the fibro-dominated HNSCC TME model with different immune accessibility indices. We observed the ICI dose response deteriorates linearly with the reduction of immune accessibility. Further, the balance between the anti- and pro-tumor roles of the T cell states modulates the slope of the dose efficacy-immune accessibility curve (Fig. 6(b)). Further, unlike the immune-dominated scenario, the post-ICI CAF population remains almost identical to the pre-ICI counterpart in the low immune-accessible scenario (Fig. 6(d)). The invariance of the CAF population to ICI in the fibro-dominated setting is one of the most critical prognostic differences between the immune-dominated and fibro-dominated TME subtypes. A simulation study of the CAF-immune accessibility index for multiple CAF proliferation rate reveals that the immune accessibility index increases gradually (smoothly) with respect to the growth of CAF (Fig. S6(a)). However, there exists a threshold timepoint dependent on the CAF proliferation rate, beyond which the immune accessibility deteriorates drastically (Fig. S6(b)). To understand the mechanism, we studied the phase space in the LIF-tumor cell plane. Unlike the immune-dominated conditions, we observed that lower immune accessibility does not significantly change the LIF levels---a significant player in the conversion from the wild-type fibroblasts to CAFs. Thereby retaining the pre-ICI conversion flux (Figure 6(c)).

As mentioned, the post-ICI responses for parametric conditions resembling the fibro-dominated and the immune-desert scenario are similar (barring T cell counts inside the whole TME) regarding the change in the tumor cells and pro-tumor components. These results motivate us to ask whether both the TME subtypes (immune-desert and fibro-dominated) require similar combinational therapies to improve the ICI response.

### Leveraging the molecular landscape of the TME-possible targets and biomarkers

According to the proposed model, the post-ICI outcomes for different pre-ICI TME subtypes are significantly distinct. Although the fibro desert and immune-dominated result in the shrinkage of the total tumor cell population, the immune-desert, and fibro-dominated TME subtypes remain largely insensitive to ICI-based treatment. Therefore, to circumvent the ICI insensitivity emergent in some TME subtypes, we explore the prospect of modulating the well-known molecular species in the HNSCC TME. The no-show of ICI response for immune-desert TMEs can be attributed to the zero (or very few) pre-ICI killer T cell population. Therefore, any combination therapy in an immune desert scenario should aim to modulate the proliferation to death and conversion balance for the killer T cell (ζ_*KillerT cell*_). For this purpose, we chose IL-2 to enhance the killer T cell population in an immune desert scenario.

On the other hand, the insensitivity to ICI in the fibro-dominated parameter regions emerges due to either a high CAF-tumor cell paracrine rate or a low immune accessibility index. The common bottleneck in both scenarios is the abundance of the CAF population. Therefore, targeting the CAF via selected molecular species may improve immune accessibility and reduce the CAF-tumor paracrine rate. For this purpose, we selected OPN and LIF as the target molecular species for both molecules, which play a crucial role in the proliferation of- and conversion to CAF.

### ICI+IL-2, reprograms the immune-desert TME

As discussed in the preceding sections, a significant fraction (due to low Killer T cell proliferation) of the immune-desert TME subtypes do not respond to ICI treatment. The primary reason is the depletion of killer PD1− killer T cells under pre-ICI conditions. Further, due to the small killer T cell population, the final, pre-ICI, IL-2 concentration settles to an extremely low level. According to the HNSCC TME network, the population of the killer T cells can be enhanced through IL-2. Therefore, we simulated an immune-desert TME with one-time, initial IL-2 spikes of varying amplitude. Interestingly, we observed that for one time, IL-2 spikes beyond a threshold level, the cell state population trajectories move away from the immune-desert to an immune-non-desert trajectory (Fig. 7(a-c)), which renders a favorable post-ICI outcome for a high immune accessibility index. The threshold one time IL-2 levels required to drive the TME away from immune desert is dependent on the balance between the IL2-driven autocrine and exhaustion, death rate of killer T cells (Fig. S7).

**Fig. 7.**
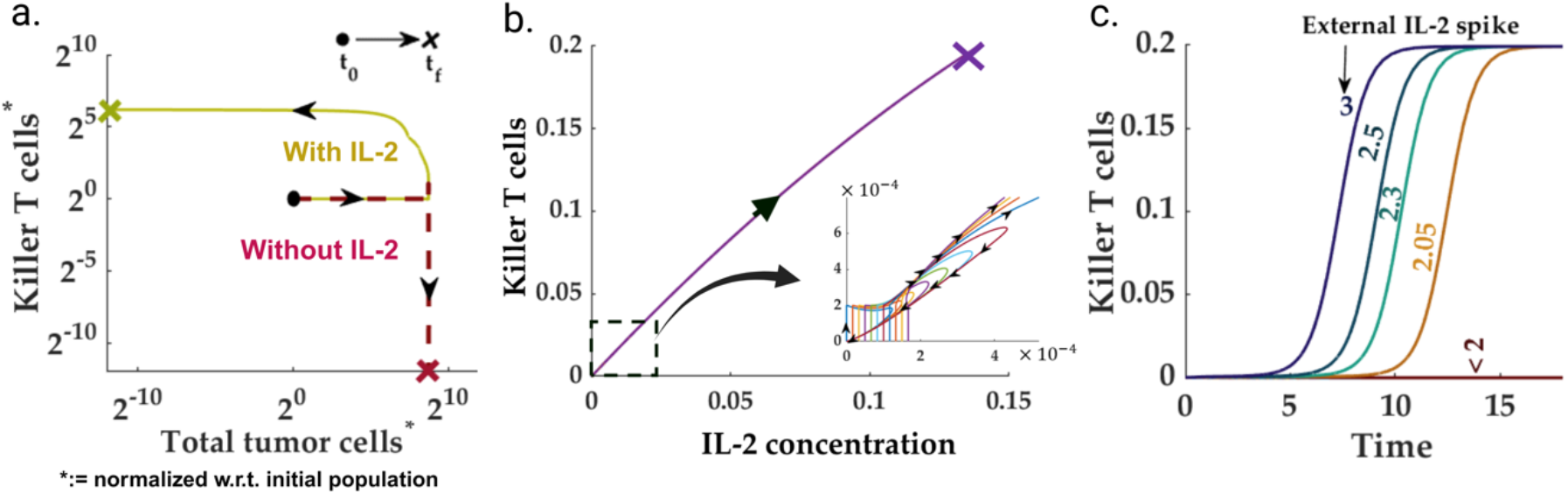
One-time IL-2 spikes— reprogramming immune-desert. **(a)** A spike in the initial IL-2 levels (beyond a threshold) can drive the system trajectories towards an immune- non-dessert steady state. **(b)** Although some trajectories return to the immune-desert arrangement, the trajectory settles in a non-zero killer T cell population beyond a threshold IL-2 injection. **(c)** Time profiles for killer T cells for different external IL-2 levels.

### Restoring sensitivity to ICI for immune-inaccessible, fibro-dominated TME

According to the proposed network, the cancer-associated fibroblasts grow through i) self-proliferation via OPN-mediated autocrine interaction and ii) LIF-induced conversion from wild-type fibroblasts. Therefore, we study the effect of OPN and LIF reduction on the ICI outcome for a fibro-dominated TME with very low immune-accessibility. To understand how the balance between the immune-accessible and immune-accessible tumor cells depends on the OPN level, we simulated the proposed HNSCC model for different OPN reduction rates. We repeated this exercise in the presence of LIF and with LIF being knocked out. We observe that the reduction of OPN modifies the balance between the tumor cells vis-à-vis immune accessibility towards the accessible tumor cells. Further, a LIF knockout and OPN reduction significantly improved over both the OPN-based treatment (Fig 8(a-b)) and LIF-knockout (Fig. S9). Interestingly, the tumor cell balance reduces drastically beyond a parameter-dependent threshold anti-OPN concentration. Moreover, the reduction in the total CAF count leads to an improvement in immune accessibility, thereby bettering the chances of recovery (Fig. 8(c)).

**Fig. 8.**
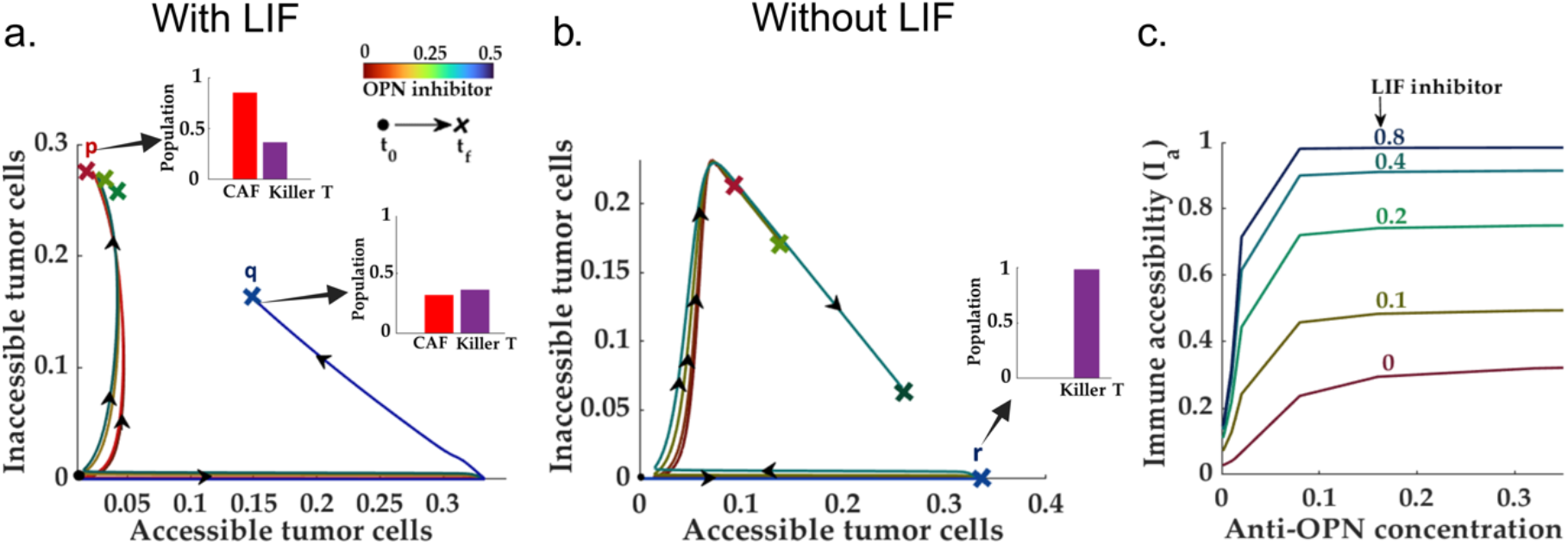
OPN and LIF knockout, from fibro-dominated to fibro-dessert. **(a-b)** Demonstrate the effect of OPN reduction on the proportion of the inaccessible to accessible tumor cells in two scenarios: with and without LIF. As shown in both cases, below a certain threshold of OPN concentration, the balance between the accessible and inaccessible tumor cells improves significantly. An additional knockout of LIF extends the OPN-knockout-driven reduction in tumor cells towards complete removal. **(c)** Suggests that OPN reduction modulates the immune accessibility index of the TME. Further, a LIF+OPN reduction drives the TME towards full accessibility.

The potential combination therapies for immune-desert and fibro-dominated are clearly different. Although the HNSCC TME subtypes produce similar diagnoses (barring the total T cell counts in the TME) vis-à-vis different tumor cell state populations, the IL-2-based treatment in the immune desert focuses on strengthening the immune system. In contrast, the OPN+LIF knockouts target the CAF population in the TME to better the immune accessibility landscape.

### IL-8: Primary distinguishing factor for different TME subtypes

As established in the preceding sections, TME subtypes play a governing role in determining the post-ICI outcomes. Unlike other cellular species, most molecular species do not exhibit specific signatures unique to TME subtypes. For instance, the IL-2 levels remain similar in both the immune-dominated and fibro-dominated scenario for the resident killer T cell population, which does not present significant variation between immune-dominated and fibro-dominated subtypes (Fig. 2(b-d)). The fold change of the OPN levels (pre-vs. post-ICI) remains indistinguishable for fibro-dominated and immune-dessert scenarios for the post-ICI CAF population. It does not undergo significant changes in the TME subtypes (Fig. 6(c)). The scenario for LIF is similar (Fig. 6(d)). Therefore, we set out to investigate the prospect of IL-8 as a potential biomarker for TME subtypes.

Interestingly, we found out that IL-8 exhibits distinct trajectories for different TME subtypes (Figures 9(a-b)). In the immune dessert scenario, due to the absence of effective killer T cells, the post-ICI IL-8 does not change. Although the immune/non-fibrotic and immune-dominated TME subtypes are marked by a significant decrease in the IL-8 levels, the absolute levels of IL-8 concentration are significantly different for both scenarios. More importantly, unlike other inflammatory cytokines such as OPN and LIF, the IL-8 level shows a slow *increase* in the post-ICI scenario for an immune inaccessible fibro-dominated scenario. This can be verified by the recent experimental observations by Hill *et al*. (2023) in HPV-negative HNSCC patients. The IL-8 levels in the plasma slightly increase for non-responsive HPV-negative patients compared to its pre-ICI counterpart.

**Fig. 9.**
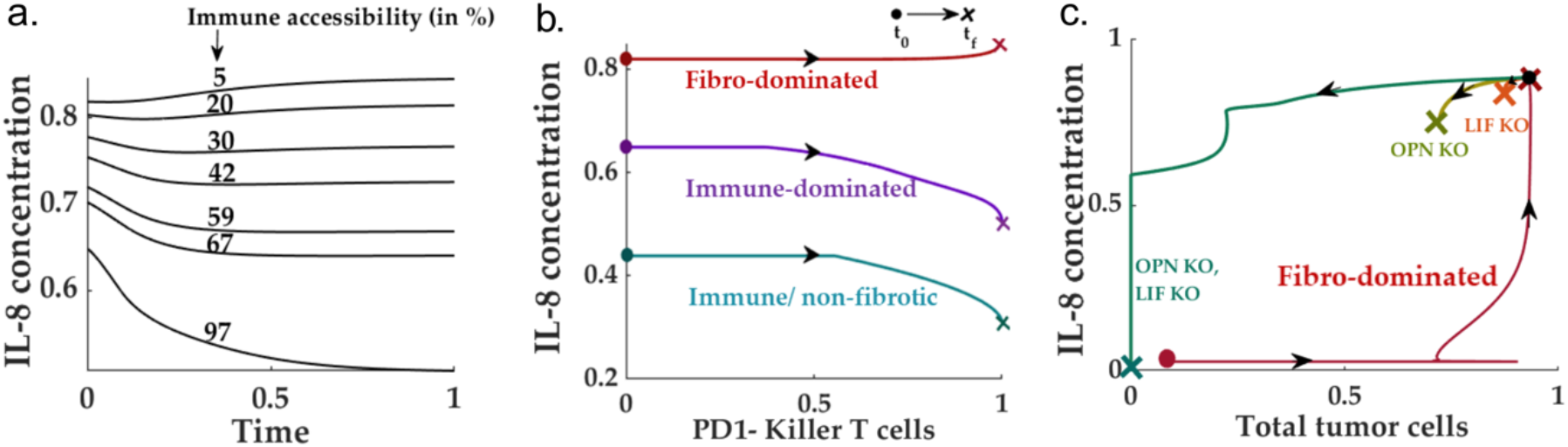
IL-8: Potential biomarker for identifying TME subtype. **(a-b)** Show that with high immune accessibility, the IL-8 level is low and is further reduced following a neo-adjuvant ICI therapy, whereas the scenario with low immune accessibility (characteristic of a fibro- dominated TME) can be identified with a slight increase in the post-ICI IL-8 level. Although the IL-8 trend is similar for immune-dominated and immune/non-fibrotic TME, the absolute levels differ significantly between the two subtypes. **(c)** The removal of both OPN and LIF reprograms the TME towards a immune/non-fibrotic subtype. Therefore, the IL-8 level post-OPN and LIF knockout followed by ICI also drops drastically compared to its pre-ICI and pre-removal counterparts.

### Lactate reduction improves the overall ICI response

Lactate is a well-known metabolite reported in the literature on several solid tumors, including HNSCC. Interestingly, a non-responsive TME has been identified with elevated lactate levels, whereas a good prognosis, in some scenarios, has been identified with lower lactate levels (91).

We leverage the proposed model to understand the effects of lactate on the population of tumor and non-tumor cells. For this purpose, given a specific set of parameters, we calculated the proportion of PDL1-to PDL1+ tumor cells for varying killer T cell cytotoxicity levels (Fig. 10(a)). We repeated the simulation for various residual lactate levels. We observed that an increasing amount of lactate pushes the TME away from immune-dominated to an effective immune-desert (or CAF-dominated) scenario (Fig. 10(b)). Therefore, targeting lactate can increase the killer T cell efficiency and move the TME to an immune-dominated region for moderate to high immune accessibility.

**Fig. 10.**
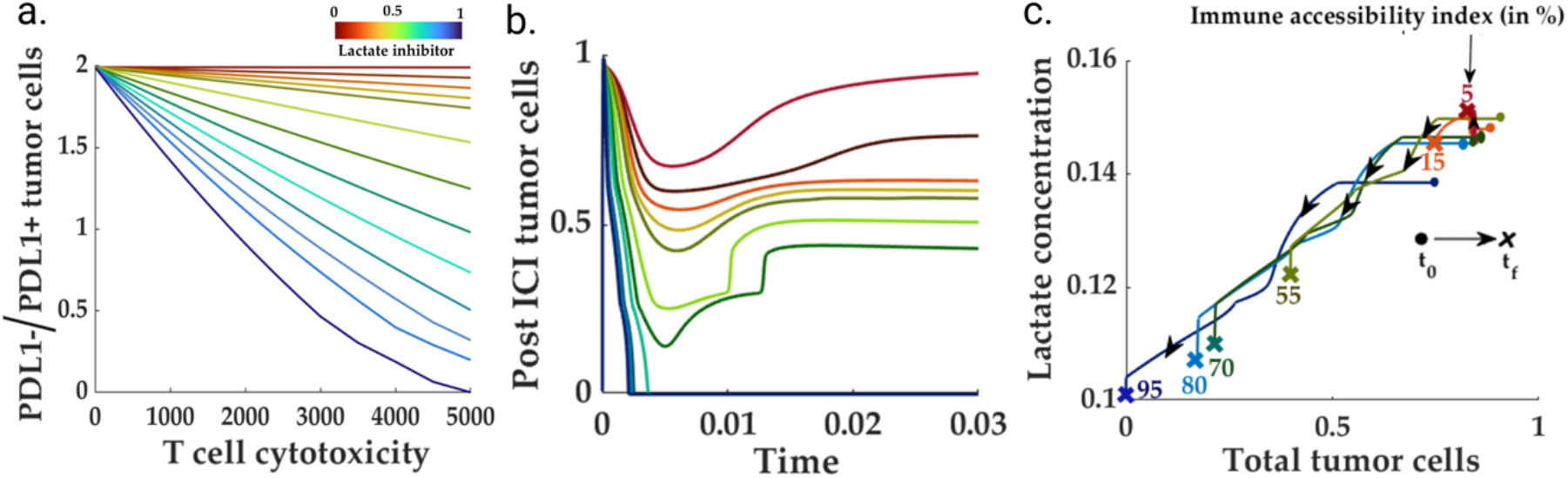
Lactate removal improves overall immune response. **(a)** Lower levels of residual lactate concentration inside the TME enhance the cytotoxicity of the killer T cells in the pre-ICI setting. **(b)** The negative impact of lactate on the immune response continues to the post-ICI scenario, wherein a high lactate level impedes the PD1-killer T cell activity, leading to a high tumor cell population. **(c)** For high-killer T cell cytotoxicity, lactate can also serve as a distinguishing factor between immune-accessible and inaccessible HNSCC TMEs. In highly immune-accessible TMEs, the application of anti- PD1 renders a significant decrease in lactate compared to its pre-ICI counterpart. In contrast, the low immune-accessible TME correlates with higher post-ICI lactate levels.

On the other hand, to study the lactate signature for different immune accessibility scenarios, we simulated the system for various values of immune accessibility. We chose the model parameters so that high immune accessibility resembles an immune-dominated scenario. We obtained the phase space for post-ICI lactate and tumor cell populations. Interestingly, the ICI intervention reduces the residual lactate levels at most immune accessibility, barring extremely inaccessible situations (Figure 10(c)). Interestingly, the post-ICI lactate level slightly increases from its pre-ICI counterpart. This suggests the potential use of lactate as a biomarker for low immune accessibility.

## Discussion

In this work, we developed a mathematical model for the HNSCC TME that explains a range of clinically (and experimentally) observed phenomena ranging from multiple TME subtypes to distinct responses to immunotherapy. Considering the trade-off between the ease of computation (theoretical/ computational analysis) and the granularity (explainability), we chose different transcriptomic cell states (and molecular species) as the agents of the proposed model. It enabled us to work with a more minor dimensional system (compared to gene regulatory network formalism) and without significant compromise on explainability. With an exhaustive search across the parameter space, we identified five distinct TME compositions, each representing clinically observed TME subtypes: fibro-dominated, immune-dominated, immune/non-fibrotic, immune-desert, and desert (4,91,92). Our model suggests that different parameter balances determine the possible path of the cell state population trajectories toward a particular TME phenotype. Whether the exact factors that govern these balances may be epigenetic or owe their emergence to mutation remains an open area of study. We observe these pre-ICI TME compositions govern the post-ICI response. Our model predicts that while the immune/non-fibrotic and immune-dominated HNSCC subtypes constitute a favorable situation for ICI therapy, immune-desert and fibro-dominated TMEs remain insensitive to therapy resistance for a wide range of model parameters. We predict the TME-specific molecular species that can modulate the ICI response for the ICI-insensitive subtypes.

Further, our model predicts multiple compositional possibilities within the immune-desert subtype due to the different mechanisms behind immune depletion. An exhaustion-driven immune desert leads to the colonization of the immune landscape by exhausted and regulatory T cells. Similar observations have been reported in the recent work by Bao *et al. (2024)*, wherein a spatial transcriptomic analysis of the immune module for twenty-six colorectal cancer patients revealed the non-responder to chemotherapy can be attributed to the immune-cold condition, defined as the abundance of regulatory T, and exhausted CD8+ and CD4+ T cells (93). Leveraging the integrative framework, our model predicts that the CAF-promoted immune cold scenario can lead to immune depletion due to the effective shutdown of the helper T cells, and the immune landscape can be hijacked by the regulatory T cells. A high exhaustion rate can also render an immune scenario colonized primarily by the exhausted and regulatory T cells.

Our analysis shows that these identified TME compositions in pre-ICI settings govern the prognosis during and after ICI therapy. Our simulation studies indicate that two pre-ICI TME compositions, resembling 1.) the immune desert and 2) fibro-dominated TME subtypes remain primarily insensitive to ICI therapy. While the ICI resistance due to the immune-desert phenotype is well established, the recent work by Quek *et al*. (2024), from the single cell analysis of five melanoma patients, showed the existence of resistant TME subtypes with abundant CAF and non-zero killer T cell population—This supports our result depicting a fibro-dominated scenario being insensitive to ICI therapy (94). Lack of clinical response in several HNSCC ICI clinical studies can be mapped to the immune-desert nature of the underlying TME. The post-ICI tumor cell count remains significant despite a meager increase in the proportion of CD8+ killers to exhausted T cells (97-98). According to our model, although an ICI intervention increases the proportion of killers to exhausted T cells, the post-ICI tumor cells remain significant due to the oncogenic role of the exhausted T cells. Our model predicts a reduction of elevated lactate levels may improve the scenario by increasing the effective cytotoxicity of the killer T cells, thereby leading to an aggressive immune response.

Interestingly, according to our model, the abundant CAF population can also lead to a condition wherein the TME, despite a significant killer T cell population, exhibits an immune-desert scenario vis-a-vis the ICI response. Recent work by Xiao *et al*. (2023) on the effect of CAF on the CAR-T cell therapy for desmoplastic pancreatic tumor patients revealed that the resident CAF population inhibits the killer T cell-driven immune response via the construction of a physical barrier between the T and Tumor cells (98). The immuno-suppressive role of CAFs and the associated pathways have also been reported in the literature (95,97,98). Bao *et al*. (2024) observed a class of non-respondent (to immunotherapy) melanoma tumors with a non-zero killer T cell population (93). Our model predicts that beyond a threshold population levels of CAF, the immune response deteriorates drastically. Further, the CAF-driven barrier divides the local tumor cells into immune-accessible and inaccessible classes. Further, due to resource competition, the application of anti-PD1 increases inaccessible immune cells, rendering the total tumor cell count largely unaffected.

On the other hand, under the parametric conditions resulting in immune/non-fibrotic and immune-dominated TME subtypes, the proposed model yields preferable outcomes to ICI. As suggested by the model, unlike the immune/non-fibrotic subtype, the pre-ICI diagnosis of an immune-dominated TME can potentially present with abundant CAFs and killer T cells. The central differentiating feature between the pre-ICI conditions for fibro-dominated and immune-dominated subtypes is the presence of CAF-protected, immune-inaccessible tumor cells. *In this sense, it may be hypothesized that fibro-dominated is an advanced scenario of immune-dominated TME*.

The model-guided approach enables us to propose possible molecular targets to circumvent the poor prognosis caused by the pre-ICI stable TME compositions. Our study shows that non-responsive outcomes to ICI treatment due to an immune-desert TME may be circumvented by a one-time IL-2 spike, which reprograms the TME towards an immune-hot scenario. Multiple studies on different cancer types, including melanoma, reported that the IL-2-based treatments (99, 100), in some scenarios, lead to a complete or partial response (99). Our model predicts that a one-time IL-2 spike beyond a threshold can drive the immune desert HNSCC TME toward an immune-hot scenario. According to our model, immune-hot TME can only yield a desirable prognosis in immune-accessible tumors, *i*.*e*., if the TME composition is not fibro-dominated. This can explain the scenario where an IL-2-based combination therapy does not yield a favorable response. Although the history of IL-2 based treatment dates back to the some of the earliest immune based therapies and carried extensive toxicity, new strategies of delivery, dosing and timing could be considered to shift a TME rather than use to drive the entire therapeutic approach (101).

Unlike the immune-desert TME, the model proposes an OPN+LIF-based intervention for the fibro-dominated scenario. As observed in this study, the OPN and LIF knockout significantly improves the accessibility landscape via a drastic reduction of the CAF population inside the TME. This is also supported by the experimental observations of elevated OPN and LIF levels in ICI-resistant patients in across different cancers (102-106). Therefore, the two non-responsive TME compositions, namely the fibro-dominated and the immune-desert, require different interventions (apart from the standard ICI) to improve the prognosis. Therefore, it is essential to differentiate these two different TME compositions. Our model proposes that IL-8, unlike most molecular species, may have distinct features for the TME subtypes. In a recent study on the response of HNSCC patients to ICI across HPV signatures, Hill *et al*. (2023) reported that the HPV-negative HNSCC patients presented with increased levels of IL8 (compared to pre-neoadjuvant ICI setting) in the scenario of no-response. Meanwhile, the post-ICI IL-8 levels significantly reduced the number of responders (88). Our model predicts that the post-ICI IL-8 remains high and identical to the pre-ICI counterpart in an immune-desert scenario due to the lack of significant change in the tumor cell and M2 macrophage population.

In contrast, the post-ICI IL-8 levels increase for a fibro-dominated scenario compared to its pre-ICI counterpart. This can be reasoned as follows: The post-ICI scenario for fibro-dominated and non-immune desert HNSCC TME can be identified with a significant (marginal) increase in killer T cells, which in turn leads to an increase in the IL-10 levels. Further, elevated IL-10 levels result in aggressive conversion to M2 macrophages from the resident M1 macrophages. Therefore, the overall population of all the IL-8 contributors in the HNSCC TME, the CAF, tumor cells, and M2 macrophages shows a marginal increase compared to its post-ICI counterpart.

The proposed model is limited by its scope and granularity and, the objective of this work. The crucial notion of *staging and metastasis* of the primary tumor over time has yet to be a part of this model. This can inspire further modification and complexity over and above the existing model schema. Moreover, like any mathematical modeling exercise, the correctness of the conclusions drawn from the proposed model is conditioned on the specific dynamic and the assumptions thereof. Throughout this work, the proposed modeling and analysis framework assumes spatial homogeneity, at least for the molecular species. This implies that the diffusion current of the molecular species is faster than the timescales of interactions. Further, since the model is built from the existing literature, there may exist a degree of confusion about the exactness of the proposed HNSCC TME network. Additionally, although the proposed model attempts to provide a comprehensive picture of the TME, it is not a complete and extensive description of TME, leaving the scope for further modification and updating of the current version.

Overall, the proposed model is able to explain a wide range of clinical observations across patients and contains the scope to be tuned to particular patient-specific quantitative information, including the single-cell transcriptome analysis and immunohistochemistry of the TME. Therefore, using the patient-specific single-cell transcriptomic dataset to tune the model can be an exciting area of future study.

## Materials and Methods

In this section, we outline the workflow of the manuscript. We first reconstructed the TME network structure for HNSCC from the literature. Subsequently, we used rate laws applicable to each of the interaction types to formulate a quantitative systems model of HNSCC TME.

### Reconstruction of the TME components and interactions from literature

We include four cell types as the constituting modules of the TME, namely i) tumor cells, 2) T cells, 3) macrophages, and 4) fibroblasts (3, 9, 32, 33). We also consider eight molecular species secreted from one or many of the four cell types. Interleukin-2 (IL-2), interleukin-6 (IL-6), interleukin-8 (IL-8), interferon gamma (IFN*γ*), lactate (Lac), intercellular adhesion molecule (ICAM), Osteopontin (OPN), and interferon regulatory factor-8 (IRF-8)— these eight molecular species are crucial for the conversion, proliferation, and apoptosis of the cell types present in the TME.

### Tumor cells

Characteristic to the TME, tumor cells are associated with high epithelial scores and high mutational burden. Recent single-cell transcriptome analysis revealed the diversity within the tumor cells. In the scenario of HNSCC, Puram *et al*. (2017) proposed seven distinct tumor cell states with diverse groups of biomarkers (30). The transcriptome analysis of a number of cancer cell lines reveals a specific stem-like pattern which, upon further analysis has been reported to transition into different tumor cell states to facilitate immune escape, metastasis, and drug resistance (3, 5, 18, 26, 28)— this may also be understood as the phenomena of ‘cooperation’ between tumor cells (34).

Since this work is centered around immuno-therapy, we classify the tumor cells based on their response to the activities of the immune cells. It is well-known that the natural killer (NK)-cells and the killer T cells recognize the tumor cells without programmed death ligand1 (PDL1) (22, 35). Interestingly, several studies confirmed that a PD-L1 expression in tumor cells is a known strategy for immune escape (22, 36, 37).

Therefore, the proposed formalism consists of three different tumor cell states— 1) tumor stem cells, 2) tumor cells without PDL1, and 3) tumor cells with PDL1 expression. Further, the tumor cells secrete inflammatory cytokines (such as OPN, IL–8) and lactate that aid in self–proliferation and sustenance in the presence of immune onslaught (38–41).

### T-cells

The T-cells, as the main driver of the immune response, destroy the stem-like and PDL1− tumor cells. The killer and helper T-cells also secrete many pro-immune cytokines such as IFNγ, IL-2, and ICAM1 (42–44). Upon a classification exercise of the transcriptome of the non-tumor cells, Puram et al. (2017), identified four different cell states within the T-cell cluster— namely 1) PD1+ killer, 2) helper T cells, 3) regulatory T cells, and 4) exhausted T cells (30).

The helper T cells promote the growth of the killer T cells in the presence of PDL1− tumor cells (45). On the other hand, regulatory T-cells control the population of helper T cells to circumvent the overactive immune response (31). Further, there also exists a significant population of killer-like T cells (also known as exhausted T-cells) that are derived from the interactions between the PD1+ killer T cells, PDL1+ tumor cells, and M2-phase macrophages. With ICI-based therapies, the killer PD1+ T cells are converted to PD1−, thereby reducing the conversion flux towards the exhausted T cells and improving the overall immune response (30, 46).

### Macrophages

Macrophages play a dual role within the TME. The M1 phase macrophages aid in the production of killer T cells (22, 47). Further, the M1 macrophages also secrete transcription factors that can play a crucial role in inhibiting the secretion of pro-inflammatory cytokines. The M1 macrophages also secrete IRF8, inhibiting OPN secretion in TME, thereby improving the prospect of ICI therapy.

Unlike the M1 phase, the M2 phase macrophages serve a pro-tumor role in several ways, such as accelerating the conversion of killer T cells to an exhausted state (48, 49).

### Fibroblasts

Fibroblasts are the most common unit apart from the tumor cells to be found inside the TME. Broadly classified in two distinct phenotypes– myofibroblasts and invasive, fibroblasts modulate the proliferation flux of the tumor cells via paracrine interactions. Further, the invasive fibroblasts can also protect the tumor cells from being attacked by the killer T cells (50, 51). Additionally, fibroblasts also secrete several cytokines (IL-8 and OPN) that can further the growth of tumor cells and hamper the immune response (52, 53).

### Modeling of the TME dynamics

This section presents the specific interactions and crosstalk between different constitutive units discussed in the preceding sections. We establish the general modeling rules and assumptions adopted throughout this work.

#### *A*.1 Rules

The following are the general rules adopted for the model.

1. The dynamics is modeled using ordinary differential equations (ODE) i.e., the cell-state populations and the molecular concentrations are modeled as functions of time.
2. Proliferation: Each cell state (x_j_) has its own proliferation flux expressed in terms of logistic growth

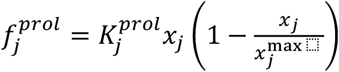

where, 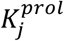, and 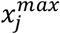 are the associated rate for proliferation and the carrying capacity of x_j_ respectively.
3. Conversion: Each cell state (x_j_) can convert to a different cell state (x_i_) within the same cell type with
4. the proposed flux rate

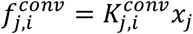

where, 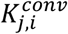 is the associated conversion rate.
5. Paracrine interaction: If a cell-state x_i_ promotes the proliferation rate of cell xj then the modified proliferation rate for x_j_ can be expressed as

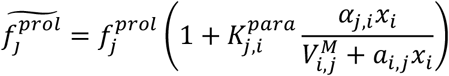

where, 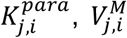, and *α*_*i,j*_are the rate constant, dissociation constant, and the spatial proximity index between x_i_ and x_j_ respectively.
6. Regulatory inhibition: The regulatory inhibition between two species x_i_ (source) and x_j_ (target) modulates the proliferation in the following manner

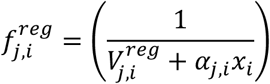

where,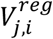, is the dissociation constant for the regulatory inhibition.
7. Elimination: If a cell state x_i_ eliminates/ destroys another cell state x_j_ the associated flux can be expressed as

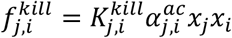

where, 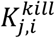 is the rate constant corresponding to the elimination reaction. The term 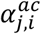 modulates the accessibility (proximity) from the cell state *x*_*i*_ to *x*_*j*_.
8. Death/Degradation: Every species is assumed to have a natural death (for cell states) or degradation (molecular species) rate that is reflected in the following manner

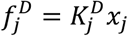

where,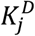 is the natural death/ degradation rate.

#### F.2. Calculating the proliferation to death balance (*ζ*)

As illustrated from our study, the clinically observed TME subtypes can be mapped as the pre-ICI steady state of the proposed TME model for HNSCC. For this purpose, we introduced a quantity ζ that is defined as the ratio between the maximum proliferation factor of a given cell state to the corresponding death and conversion rate. We calculate the maximum proliferation factor 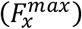 for a given cell state (x) in the following manner:

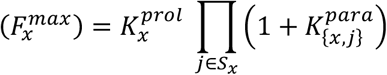

Where, *S*_*x*_ is the set of all the incoming, neighbor nodes of x. It is to be noted that since the paracrine interaction dynamics, according to the proposed model, is rational,polynomial bounded above by the paracrine rate constant the net proliferation rate of a cell state is always bounded above by 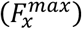.

On the other hand, the depreciation of the cell state x occurs through the processes of killing, conversion, and natural death. Therefore, the log-transformed net proliferation to death balance ζ can be calculated as

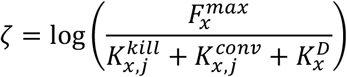

Now, for total tumor cells, we used the maximum ζ across different tumor cell states.

### CAF-mediated alteration of immune accessibility

It is well-known in literature that CAF plays a crucial role in remodeling the extra cellular matrix and thereby altering the accessibility from the T cells to the tumor cells. To address this critical phenomenon in the proposed model, we defined the quantity immune accessibility index (0 ≤ *I* _*a*_≤ 1) -a bounded hyper-parameter defined as the ratio between the total carrying capacity of immune accessible tumor cells to that of fibroblast protected tumor cells. We also proposed an expression for the immune accessibility index. We observed that the lack of immune accessibility alters the TME towards a fibro-dominated environment. We propose the following way to calculate immune accessibility index (I_a_)

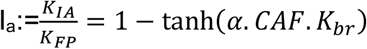

where, *K*_*IA*_ and *K*_*FP*_ denote the carrying capacities of immune-accessible and CAF-protected tumor cells respectively, α determines the fraction of CAFs near the tumor cells, and *K*_*br*_ is the barrier formation rate for the CAFs. As depicted in the expression for the immune accessibility index, is a bounded quantity in the closed interval [0,1] wherein I_a_=1 refers to the situation where all the tumor cells are accessible to the immune system, whereas I_a_=0 denotes the impenetrability of the immune cells to the TME.

As established before, the CAF population around the tumor cell states significantly affects immune accessibility. Therefore, for any given CAF population, α, and *K*_*br*_ we propose the following rule for the penetration of the immune cells 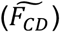 as

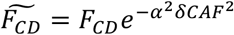

where δ is the width of CAF barrier.

### Simulation

All the simulations are done using MATLAB 2022B (Mathworks Inc., Natick, MA). The ordinary differential equations in the models are simulated using ode23s in MATLAB 2022B. The figures have been created using Biorender.com.

## Supporting information

Supplemental figures, tables and model description

## Acknowledgments

The authors thank Alexandra Machel for reviewing the MATLAB code and replicating the figures of the main manuscript. The authors are thankful to Prem Jagadeesan for independently implementing the model in MATLAB based on the manuscript text and supplement to reproduce the Figure 3 of the main manuscript.

## Code availability

The necessary MATLAB code and data required to reproduce the figures can be accessed at https://github.com/Daniel-Baugh-Institute/HNSCC_TME_Model

## Figure Legends

**Figure 1. Construction of cell-state specific tumor-microenvironment network for HNSCC:** We begin with the relevant cell types in an HNSCC TME obtained from the literature. The existing cell-type-specific models do not capture the interactions in a reliable manner for there may exist mutually opposite effects between two cell types (*e*.*g*., T cells and tumor cells). Subsequently, the literature on single-cell studies of HNSCC TME reveals the functionally distinct cellular states for different cell types (Puram *et al*. (2019)). Similarly, we searched for the possible interactions between different cell states of interest to establish a cell-state interaction network. Further, we considered relevant molecular species from the HNSCC literature that affect the population of individual cell states.

**Figure 2. Explaining the phenotypic heterogeneity of the HNSCC TME. (a)** The nodes are either the cell states or the molecular species, whereas the edges represent diverse forms of interactions. The acronyms C_0, C_PDL1+, and C_PDL1- refer to stem, PDL1+ (programed death ligand1), and PDL1- tumor cells, respectively. T_K+, T_K-, T_Help, T_Reg, and T_Ex stands for PD1+ (programmed death 1), PD1- killer T cells, Helper T cells, Regulatory T cells, and Exhausted T cells, respectively. M_1 and M_2 refer to macrophages of M1 and M2 phase, respectively. Further, F_WT and CAF correspond to wild type and invasive cancer associated fibroblasts, respectively. The acronyms IL-2, IL- 8, LIF, IFNG, IRF8, OPN, ICAM1, and Lac denote Interleukin-2, Interleukin-8, Leukemia Inhibitory Factor, Interferon Gamma, Interferon Regulatory Factor-8, Osteopontin, Intercellular Adhesion Molecule-1, and Lactate, respectively. **(b)** Based on the reconstructed network system, we simulated the dynamics for 10,000 different choices of parameters to obtain the possible steady state clusters. We found five distinct clusters namely desert, immune desert, fibro desert, immune-dominated, and fibro dominated. **(c-g)** Bar charts for the median population of tumor cells, killer T cells, CAF population across different cohorts. The lower and upper intervals indicate the 25% and 75% quartiles, respectively.

**Figure 3. *The immune-desert scenario:* (a-b)** Demonstrate a high T-cell exhaustion program that can drive an immune system with a high proliferation rate to the immune- desert phenotype. **(c)** Demonstrates the possibility of an immune-cold scenario with abundant regulatory T cells due to high CAF-Treg interaction. **(d)-(e)** Unlike other immune-desert phenotypes, the ICI intervention can improve the exhaustion-driven immune desert scenario. **(f)** Lactate reduction can further reduce the total post-ICI tumor cell count.

**Figure 4. *Immune/non-fibrotic, favorable case, and recurrence:* (a-b)** Shows despite the absence of CAF, the pre-ICI PDL1^-^ tumor cells critically depend on the balance between the oncogenic (via T_exh_) and cytotoxic activities of T cells. **(c)** The intervention of ICI drives towards an aggressive cytotoxic T cell driven immune response. **(d)** Indicates the pivotal role of residual fixed resource supply rate (nutrients, blood flow, oxygen supply) in governing the possibility of recurrence despite a favorable prognosis.

**Figure 5. *Immune-dominated-- High immune accessibility, moderate CAF:*** For a given CAF-tumor interaction rate, lower T-cell cytotoxicity leads to higher PDL1^-^ to PDL1^+^ tumor cell population. **(a)** For high levels of cytotoxicity, the tumor cell population remains insensitive of the proliferation rate () of the killer T cells in the immune dominated scenario. CAF plays a dual role in governing the helper T cell population. CAF reduces the helper T cells via the regulatory T cells. However, a high tumor-promoting role of CAF increases the resident PDL1- tumor cells that, in turn, increases the helper T cells via antigen sensing mechanism. **(c**) A high exhaustion rate can potentially increase the share of PDL1- tumor cell population owing to the abundance of the exhausted tumor cells and its tumor-promoting effects. (**d)** The post-ICI scenario significantly reduces the tumor cells and CAF. The application of anti-PD1 reduces the resident CAF population and drastically reduces the tumor cell population. (**e)** The post-ICI tumor cell population is proportional to the tumor promoting role (via paracrine interaction) of the remaining CAF population.

**Figure 6. *Fibro dominated, a modified immune desert:* (a)** depicts the conceptual framework of the proposed hypothesis on the CAF-pockets protected from the immune response. **(b)** demonstrate the fact that the immune accessibility index governs the post-ICI proportion of immune-accessible and immune-inaccessible tumor cells. **(C)** Contrary to the immune-dominated TME, the post-ICI CAF population remains almost constant at low immune accessibility due to a lack of change in the post-ICI LIF levels compared to its pre-ICI counterpart. **(d)** Phase-space for LIF and total tumor cell population. The change in LIF levels due to the ICI therapy reduces with respect to decreasing accessibility.

**Figure 7. *One time IL-2 spikes— reprogramming immune-desert*: (a)** A spike in the initial IL-2 levels (beyond a threshold) can drive the system trajectories towards an immune-non-dessert steady state. **(b)** Indicates that although some of the trajectories return to the immune-desert arrangement, the trajectory settles to a non-zero killer T cell population beyond a threshold IL-2 injection. **(c)** establishes the effect of IL-2 based treatment on the efficiency of ICI.

**Figure 8. *OPN and LIF knockout, from fibro-dominated to fibro-dessert*: (a-B)** Demonstrate the effect of OPN reduction on the proportion of the inaccessible to accessible tumor cells in two scenarios: with and without LIF. As shown in both the cases, below a certain threshold of OPN concentration, the balance between the accessible and inaccessible tumor cells improves significantly. This is further extended towards complete removal with the knockout of LIF. **(C)** Suggests that OPN reduction modulates the immune accessibility index of the TME. Further, a LIF+OPN reduction drives the TME towards complete accessibility.

**Figure 9. *IL-8: Potential biomarker for identifying TME subtype*: (a-b)** Show that with high immune accessibility, the IL-8 level is low and is further reduced following a neo- adjuvant ICI therapy, whereas the scenario with low immune accessibility (characteristic of a fibro-dominated TME) can be identified with a slight increase in the post-ICI IL-8 level. Although the IL-8 trend is similar for immune-dominated and immune/non-fibrotic TME the absolute levels differ significantly between the two subtypes. **(c)** Illustrates the role of OPN and LIF knockout on the IL-8 profile. The removal of both OPN and LIF reprograms the TME towards a immune/non-fibrotic subtype. Therefore, the IL-8 level post OPN and LIF knockout followed by ICI also drops drastically compared to its pre-ICI and pre-removal counterpart.

**Figure 10. *Lactate removal improves overall immune response:* (a)** Lower levels of residual lactate concentration inside the TME improves the cytotoxicity of the killer T cells in the pre-ICI setting. **(b)** The negative impact of lactate on the immune response continues to the post-ICI scenario wherein a high lactate levels impedes on the PD1^-^killer T cell activity thereby leading to a high tumor cell population. **(c)** In the presence of high killer T cell cytotoxicity, lactate can also serve as a distinguishing factor between immune accessible and inaccessible HNSCC TMEs. In the highly immune accessible TMEs, application of anti-PD1 renders significant decrease of lactate compared to its pre-ICI counterpart whereas, the low immune accessible TME corelates with higher post-ICI lactate levels.

