## Supplemental figures, tables and model description for "Tumor microenvironment governs the prognostic landscape of immunotherapy for head and neck squamous cell carcinoma: A computational model-guided analysis"

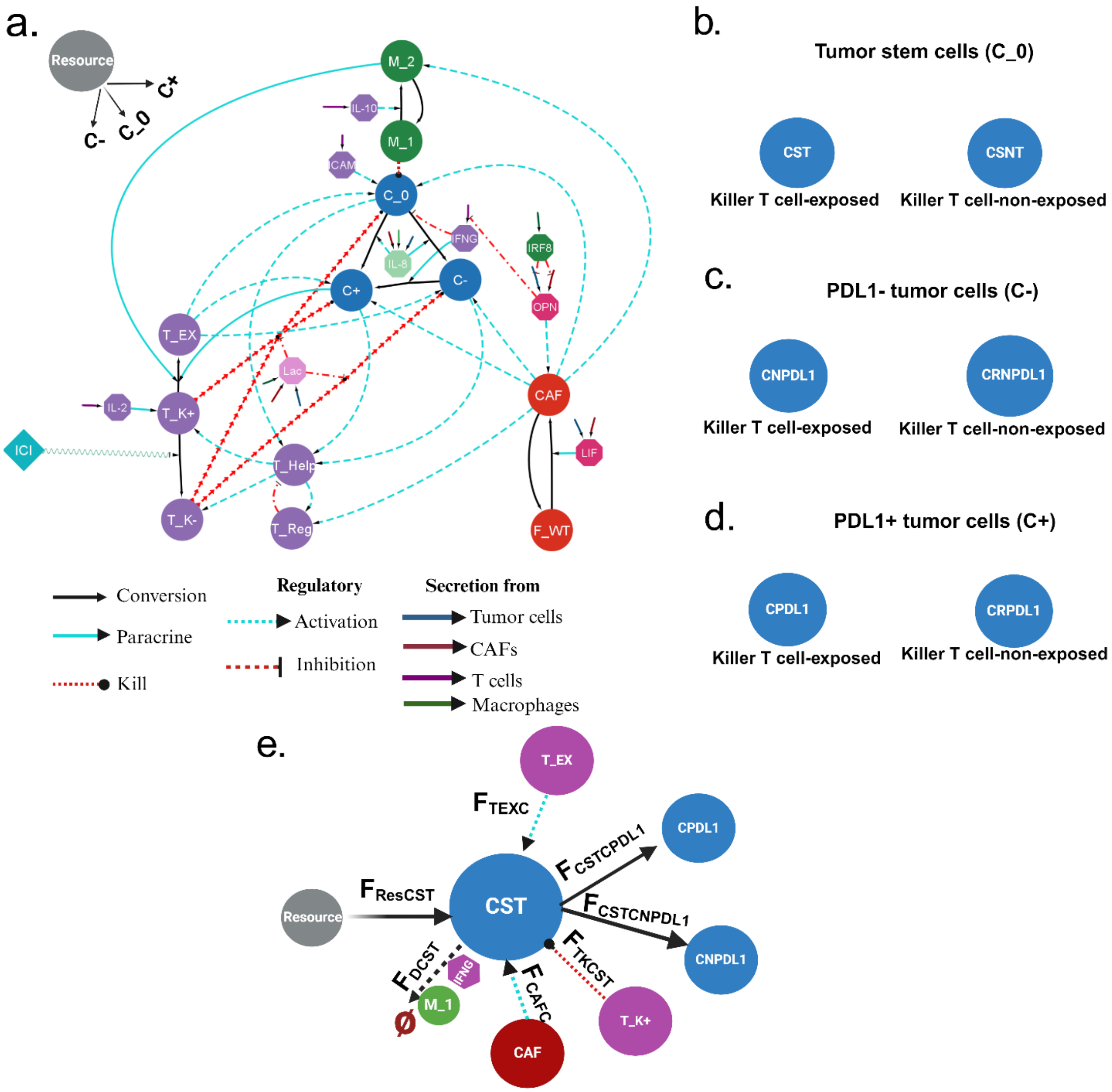


**Figure S1. Detailed HNSCC TME network model: (a)** The nodes are either the cell states or the molecular species, whereas the edges represent diverse forms of interactions. The acronyms C_0, C+, and C– refer to stem, PDL1+ (programed death ligand1), and PDL1– tumor cells, respectively. T_K+, T_K-, T_Help, T_Reg, and T_EX stands for PD1+ (programmed death 1), PD1- killer T cells, Helper T cells, Regulatory T cells, and Exhausted T cells, respectively.

M_1 and M_2 refer to macrophages of M1 and M2 phase, respectively. Further, F_WT and CAF correspond to wild type and invasive cancer associated fibroblasts, respectively. The acronyms IL-2, IL-8, IL-10 LIF, IFNG, IRF8, OPN, ICAM1, and Lac denote Interleukin 2, Interleukin 8, Interleukin 10, Leukemia Inhibitory Factor, Interferon Gamma, Interferon Regulatory Factor 8, Osteopontin, Intercellular Adhesion Molecule 1, and Lactate, respectively. All the cell states are assumed to be capable of self-proliferation and natural death. Therefore, the self-loops are not shown for brevity. **(b-d)** Each tumor cell state is subdivided depending on the accessibility from the Killer T cells. The Killer T cell-exposed tumor cells are exposed to immune response whereas the Killer T cell-non-exposed tumor cells are protected by the CAF-derived barrier from immune onslaught. **(e)** Flux-structure mapping for the killer T-cell-expose tumor stem cells.

**Table S1:** Mathematical representation of the fluxes corresponding to the HNSCC TME network in Figure S1.

| Reaction flux | Mathematical expression |
|---|---|
| Proliferation of CST ($F_{ResCST}$) | $$ResK_{RCST}CST\left(1-\frac{CST}{(1-I_a)Y_{CST}+1}\right)\alpha_C$$ $$\alpha_C \coloneqq \frac{1}{a_{comp}(CPDL1+CNPDL1)+1}$$ $$I \coloneqq \tanh\,(\alpha K_{CAFB}CAF)$$ $\alpha$: Proportion of CAF engaged in barrier formation<br>$K_{CAFB}$: Barrier formation rate |
| Exhausted T-cells-driven proliferation modulator of tumor cells ($F_{TEXC}$) | $$\left(1+K_{TXC}\frac{TEX}{TEX+1}\right)$$ |
| CAF-driven proliferation modulator of T-exposed Tumor cells ($F_{CAFC}$) | $$\left(1+K_{CAFC}\frac{CAF}{CAF+1}\right)$$ |
| Conversion from CST to CNPDL1($F_{CSTCNPDL1}$) | $$K_{CSTCNPDL1}\left(\frac{IL8}{IL8+1}\right)CST$$ |
| Conversion from CST to CPDL1($F_{CSTCPDL1}$) | $$K_{CSTCPDL1}\left(\frac{IL8}{IL8+1}\right)CST$$ |
| Killer T cell-driven elimination($F_{TKCST}$) | $$K_{TKC}CST(TKPD1+TKNPD1)\gamma\beta$$ $$\gamma \coloneqq \frac{IFNG}{IFNG+1}$$ $$\beta \coloneqq \frac{1}{K_{LAC}LAC+1}$$ |
| Death of CST ($F_{DCST}$) | $$(K_{CSTD}+\delta_{IFNGCSTD}+\delta_{M1CSTD})CST$$ $$\delta_{IFNGCSTD} \coloneqq K_{IFNGCSTD}\frac{IFNG}{IFNG+1}$$ $$\delta_{M1CSTD} \coloneqq K_{M1CSTD}\frac{MACM1}{MACM1+1}$$ |

| | |
|---|---|
| Proliferation of CSNT ($F_{RESCSNT}$) | $ResK_{RCSNT}CSNT\left(1-\frac{CSNT}{I_aY_{CST}+1}\right)\alpha_{C1}$<br>$\alpha_{C1} := \frac{1}{\alpha_{comp}(CRNPDL1+CRPDL1)+1}$ |
| Exhausted T cell-driven proliferation for immune inaccessible tumor cells ($F_{TEXCR}$) | $\left(1+\alpha_T K_{TXC}\frac{TEX}{TEX+1}\right)$<br>$\alpha_T := \exp(-\delta\alpha^2 CAF^2)$<br>$\delta$: Width of CAF barrier, $\alpha$: proportion of CAF engaged in barrier forming |
| CAF-driven proliferation for immune inaccessible tumor cells ($F_{CAFCR}$) | $\left(1+K_{CAFCR}\frac{CAF}{CAF+1}\right)$ |
| Conversion from immune inaccessible stem to immune inaccessible tumor cells ($F_{CSNTCRNPDL1}$) | $K_{CSNTCNPDL1}CSNT\frac{IL8}{IL8+1}$ |
| Conversion from immune inaccessible stem to immune inaccessible PDL1+ tumor cells ($F_{CSNTCRPDL1}$) | $K_{CSNTCPDL1}CSNT\frac{IL8}{IL8+1}$ |
| Killer T cell-driven elimination of immune inaccessible tumor stem cell ($F_{TKCSNT}$) | $K_{TKC}\alpha_T(TKPD1+TKNPD1)CSNT\gamma\beta$ |
| Death of immune-inaccessible stem cells ($F_{DCSNT}$) | $F_{CSTD}\frac{CSNT}{CST}$ |
| Resource-driven growth of PDL1-immune-accessible tumor cells ($F_{RESCNPDL1}$) | $ResK_{RCNPDL1}\gamma_{CNPDL1}CNPDL1\alpha_{C2}$<br>$\alpha_{C2} = \frac{1}{\alpha_{Comp}(CST+CPDL1)+1}$<br>$\gamma_{CNPDL1} = \left(1-\frac{CNPDL1}{(1-I_a)Y_{CNPDL1}+1}\right)$ |

| | |
|---|---|
| Killer T cell-driven elimination of PDL1- immune-accessible tumor cells ($F_{TKCNPDL1}$) | $K_{TKCNPDL1}CNPDL1(TKPD1+TKNPD1)\gamma\beta$ |
| Conversion from PDL1- to PDL1+ tumor cells ($F_{CNPDL1CPDL1}$) | $K_{CPDNPD}\dfrac{IFNG}{IFNG+1}CNPDL1$ |
| Death of PDL1-, immune-accessible tumor cells ($F_{DCNPDL1}$) | $K_{CNPDL1D}CNPDL1$ |
| Resource-driven growth of PDL1+, immune-accessible tumor cells ($F_{RESCPDL1}$) | $ResK_{RCPDL1}\gamma_{CPDL1}CPDL1\alpha_{C3}$<br>$\alpha_{C3}=\dfrac{1}{\alpha_{Comp}(CST+CNPDL1)+1}$<br>$\gamma_{CPDL1}=\left(1-\dfrac{CPDL1}{(1-I_a)Y_{CPDL1}+1}\right)$ |
| Killer T cell-driven elimination of PDL1+ immune-accessible tumor cells($F_{TKNPDCPDL1}$) | $K_{TKCPDL1}CPDL1(TKNPD1)\gamma\beta$ |
| Death of PDL1+, immune-accessible tumor cells ($F_{DCPDL1}$) | $K_{CPDL1D}CPDL1$ |
| Resource-driven growth of PDL1-, immune-inaccessible tumor cells ($F_{ResCRNPDL1}$) | $ResK_{RCRNPDL1}\gamma_{CRNPDL1}CRNPDL1\alpha_{C4}$<br>$\alpha_{C4}=\dfrac{1}{\alpha_{Comp}(CSNT+CRPDL1)+1}$<br>$\gamma_{CRNPDL1}=\left(1-\dfrac{CRNPDL1}{I_aY_{CRNPDL1}+1}\right)$ |
| Death of PDL1-, immune-inaccessible tumor cells ($F_{DCRNPDL1}$) | $K_{CNPDL1D}CRNPDL1$ |
| Killer T cell-driven elimination of immune-inaccessible, PDL1- tumor cells ($F_{TKCRNPDL1}$) | $K_{TKC}CRNPDL1T_K\gamma\beta\alpha_T$<br>$T_K\coloneqq(TKPD1+TKNPD1)$ |

| | |
|---|---|
| IFNG-induced conversion to PDL1+ immune-inaccessible Tumor cells ($F_{CRNPDL1CRPDL1}$) | $K_{CPDNPD}\frac{IFNG}{IFNG+1}CRNPDL1$ |
| Resource-driven growth of PDL1+, immune-inaccessible tumor cells ($F_{ResCRPDL1}$) | $ResK_{RCRPDL1}\gamma_{CRPDL1}CRPDL1\alpha_{C5}$<br>$\alpha_{C5}=\frac{1}{\alpha_{Comp}(CSNT+CRNPDL1)+1}$<br>$\gamma_{CRNPDL1}=\left(1-\frac{CRPDL1}{I_a Y_{CRPDL1}+1}\right)$ |
| Death of PDL1+, immune-inaccessible tumor cells ($F_{DCRPDL1}$) | $K_{CPDL1D}CRPDL1$ |
| Proliferation of Killer PD1+ T cells ($F_{\mathrm{Pro}\,TKPD1}$) | $K_{TKPD}TKPD1\left(1-\frac{TKPD1}{Y_{TKM}-TEX-TKNPD1+1}\right)$ |
| M1 macrophage, Helper-driven growth of Killer T cells ($F_{THTKPD1}$) | $1+K_{THTK}\frac{TH\,MACM1}{MACM1\,TH+1}$ |
| IL2-driven growth of Killer T cells ($F_{IL2TK}$) | $1+K_{IL2TK}\frac{IL2}{IL2+1}$ |
| Effect of anti-PD1(u): Conversion from PD1+ to PD1- killer T cell($F_{TKPD1TKNPD1}$) | $K_{TKPDNPD1}TKPD1u$<br>$u$: Anti-PD1 dosage |
| Exhaustion rate ($F_{TKPD1TEX}$) | $K_{TKPDTEX}T_{KPD1}\frac{CPDL1\,MACM2}{CPDL1\,MACM2+1}$ |
| Death rate of PD1+ killer T cell ($F_{DTKPD1}$) | $K_{TKPDD}TKPD1$ |
| Proliferation of PD1- Killer T cells($F_{\mathrm{Pro}\,TKNPD1}$) | $K_{TKNPD}TKNPD1\left(1-\frac{TKNPD1}{Y_{TKM}-TEX-TKPD1+1}\right)$ |

| | |
|---|---|
| Helper-driven growth of Killer T cells ($F_{THTKNPD1}$) | $1 + uK_{THTK}\frac{TH}{TH + 1}$ |
| Death rate of PD1-Killer T cells ($F_{DTKNPD1}$) | $K_{TKPDD}TKNPD1$ |
| Proliferation rate of helper T cells ($F_{\mathrm{Pro}\,TH}$) | $K_{TH}\left(1 - \frac{TH}{Y_{TH}}\right)$ |
| Growth via antigen sensing ($F_{CANTH}$) | $1 + K_{CANTH}\frac{(CNPDL1 + uCPDL1)}{(CNPDL1 + uCPDL1 + 1)}$ |
| Regulator-driven inhibition ($F_{TREGTH}$) | $\frac{1}{1 + K_{REGTH}TREG}$ |
| Death of helper T cells ($F_{DTH}$) | $K_{THD}TH$ |
| Proliferation of regulatory T cells ($F_{\mathrm{Pro}\,TREG}$) | $K_{TREG}TREG\left(1 - \frac{TREG}{Y_{TREG}}\right)$ |
| CAF-driven proliferation of Regulatory T cells ($F_{CAFTREG}$) | $1 + K_{CAFTREG}\frac{CAF}{CAF + 1}$ |
| Death rate of Regulatory T cells ($F_{DTREG}$) | $K_{TREGD}TREG$ |
| Proliferation of exhausted T cells ($F_{\mathrm{Pro}\,TEX}$) | $K_{TEX}TEX\left(1 - \frac{TEX}{Y_{TKM} - TKNPD1 - TKPD1 + 1}\right)$ |
| Death rate of exhausted T cells ($F_{DTEX}$) | $K_{TEXD}TEX$ |
| Proliferation of wild-type fibroblasts ($F_{\mathrm{Pro}\,FWT}$) | $K_{FWT}FWT\left(1 - \frac{FWT}{Y_{FM} - CAF + 1}\right)$ |

| | |
|---|---|
| Conversion from wild type to invasive fibroblasts ($F_{FWTCAF}$) | $K_{FWTCAF}\frac{(\alpha_{LIFFWT}\,LIF)^2}{(\alpha_{LIFFWT}\,LIF)^2+K_{LIFT}}FWT$<br>$\alpha_{LIFFWT}$: Proportion of LIF in contact with FWT |
| Death of wild type fibroblasts ($F_{DFWT}$) | $K_{FWTD}FWT$ |
| Proliferation of invasive fibroblasts ($F_{\Pr oCAF}$) | $K_{CAF}CAF\left(1-\frac{CAF}{Y_{FM}-FWT+1}\right)$ |
| OPN-induced growth of invasive fibroblasts ($F_{OPNCAF}$) | $1+K_{OPNCAF}\frac{OPN}{OPN+1}$ |
| M2 macrophage-induced growth ($F_{M2CAF}$) | $1+K_{M2CAF}\frac{MACM2}{MACM2+1}$ |
| Tumor cells-driven growth ($F_{CANCAF}$) | $1+K_{CTCAF}\left(\frac{TUM_{AC}}{TUM_{AC}+1}+K_{CTCAFR}\frac{TUM_{IAC}}{TUM_{IAC}+1}\right)$<br>$TUM_{AC}\coloneqq(CST+CPDL1+CNPDL1)$<br>$TUM_{IAC}\coloneqq(CSNT+CRPDL1+CRNPDL1)$ |
| Conversion from invasive to wild type fibroblasts ($F_{CAFFWT}$) | $K_{CAFFWT}CAF$ |
| Death of invasive fibroblasts ($F_{DCAF}$) | $K_{CAFD}CAF$ |
| Proliferation of M1-macrophages ($F_{\Pr oMACM1}$) | $K_{M1}MACM1\left(1-\frac{MACM1}{Y_{MM}-MACM2}\right)$ |
| Proliferation of M1 macrophage via antigen-sensing ($F_{CANMACM1}$) | $1+K_{CANM1}\frac{(TUM_{PDL1-}+uTUM_{PDL1+})}{(TUM_{PDL1-}+uTUM_{PDL1+}+1)}$<br>$TUM_{PDL1-}\coloneqq(CST+CSNT+CNPDL1+CRNPDL1)$<br>$TUM_{PDL1+}\coloneqq(CPDL1+CRPDL1)$ |
| Conversion from M1 to M2 macrophage ($F_{M1M2}$) | $K_{M1M2}\frac{(\alpha_{IL10M1}IL10)^2}{(\alpha_{IL10M1}IL10)^2+1}$<br>$\alpha_{IL10M1}\coloneqq$ Proportion of IL-10 in contact with MACM1 |

| | |
|---|---|
| Conversion from M2 to M1 macrophage ($F_{M2M1}$) | $K_{M2M1}MACM2$ |
| Death of M1 macrophage ($F_{DMACM1}$) | $K_{M1D}MACM1$ |
| Proliferation of M2 macrophage ($F_{\Pr oMACM2}$) | $K_{M2}MACM2\left(1-\frac{MACM2}{Y_{MM}-MACM1+1}\right)$ |
| CAF-driven growth of M2 macrophage ($F_{CAFMACM2}$) | $K_{CAFM2}\frac{CAF}{CAF+1}$ |
| Death of M2 macrophage ($F_{DMACM2}$) | $K_{M2D}MACM2$ |
| IL-2 secretion by Killer Cells ($F_{TKIL2}$) | $K_{TKIL2}(TKPD1+TKNPD1)$ |
| Degradation of IL-2 ($F_{DIL2}$) | $K_{IL2D}IL2$ |
| LIF secretion by CAF ($F_{CAFLIF}$) | $K_{CAFLIF}CAF$ |
| LIF secretion by Tumor cells ($F_{CANLIF}$) | $K_{CANLIF}(TUM_{AC}+TUM_{IAC})$ |
| Degradation of LIF ($F_{DLIF}$) | $K_{LIFD}LIF$ |
| IFNG secretion by T cells ($F_{TKIFNG}$) | $K_{TIFNG}(TKPD1+TKNPD1)$ |
| Inhibition of IFNG secretion by OPN ($F_{OPNIFNG}$) | $\frac{1}{\alpha_{OPNIFNG}OPN+1}$<br>$\alpha_{\mathrm{OPNIFNG}}$: Proportion of OPN in contact with IFNG |

| | |
|---|---|
| Degradation of IFNG ($F_{DIFNG}$) | $K_{IFNGD} IFNG$ |
| IL-8 secretion by M2 macrophage ($F_{M2IL8}$) | $K_{M2IL8} MACM2$ |
| IL8-secretion by CAF ($F_{CAFIL8}$) | $K_{CAFIL8} CAF$ |
| IL8-secretion by tumor cells ($F_{CAFIL8}$) | $K_{CAFIL8}(TUM_{AC} + TUM_{IAC})$ |
| Degradation of IL8 ($F_{DIL8}$) | $K_{IL8D} IL8$ |
| Lactate secretion by tumor cells ($F_{CANLAC}$) | $K_{CANLAC}(TUM_{AC} + TUM_{IAC})$ |
| Lactate secretion by M2 macrophage ($F_{M2LAC}$) | $K_{M2LAC} MACM2$ |
| Lactate degradation ($F_{DLAC}$) | $K_{LACD} LAC$ |
| IL10 secretion by Killer T cells ($F_{TKIL10}$) | $K_{TKIL10}(TKPD1 + TKNPD1)$ |
| Degradation of IL10 ($F_{DIL10}$) | $K_{IL10D} IL10$ |
| ICAM1 secretion by Killer T cells ($F_{TKICAM1}$) | $K_{TKICAM}(TKPD1 + TKNPD1)$ |
| ICAM1 degradation ($F_{DICAM1}$) | $K_{ICAM1} ICAM1$ |
| OPN secretion by CAF ($F_{CAFOPN}$) | $K_{CAFOPN} CAF$ |
| OPN secretion by tumor cells ($F_{CANOPN}$) | $K_{CANOPN}(TUM_{AC} + TUM_{IAC})$ |

| | |
|---|---|
| Inhibition of OPN secretion by IRF8 ($F_{IRF8OPN}$) | $\frac{1}{\alpha_{IRF8OPN}IRF8+1}$<br>$\alpha_{IRF8OPN}$: Proportion of IRF8 in contact with OPN |
| Degradation of OPN ($F_{DOPN}$) | $K_{OPND}OPN$ |
| IRF8 secretion by M1 macrophage ($F_{M1IRF8}$) | $K_{M1IRF8}MACM1$ |
| Degradation of IRF8 ($F_{DIRF8}$) | $K_{IRF8D}IRF8$ |

**Model equations:** Given the fluxes in Table 1, we construct the overall mathematical model for the HNSCC TME.

$$\frac{dCST}{dt} = F_{RESCST}F_{TEXC}F_{CAFC} - F_{CSTCNPDL1} - F_{CSTCPDL1} - F_{TKCST} - F_{DCST}$$

$$\frac{dCSNT}{dt} = F_{RESCSNT}F_{TEXCR}F_{CAFCR} - F_{CSNTCRNPDL1} - F_{CSNTCRPDL1} - F_{TKCSNT} - F_{DCSNT}$$

$$\frac{dCNPDL1}{dt} = F_{RESCNPDL1}F_{TEXC}F_{CAFC} + F_{CSTCNPDL1} - F_{CNPDL1CPDL1} - F_{TKCNPDL1} - F_{DCNPDL1}$$

$$\frac{dCPDL1}{dt} = F_{RESCPDL1}F_{TEXC}F_{CAFC} + F_{CSTCPDL1} + F_{CNPDL1CPDL1} - F_{TKNPDCPDL1} - F_{DCPDL1}$$

$$\frac{dCRNPDL1}{dt} = F_{RESCRNPDL1}F_{TEXCR}F_{CAFCR} + F_{CSNTCRNPDL1} - F_{TKCRNPDL1} - F_{CRNPDL1CRPDL1} - F_{DCRNPDL1}$$

$$\frac{dCRPDL1}{dt} = F_{RESCRPDL1}F_{TEXCR}F_{CAFCR} + F_{CSNTCRPDL1} + F_{CRNPDL1CRPDL1} - F_{TKCRPDL1} - F_{DCRPDL1}$$

$$\frac{dRES}{dt} = K_{RIN}(Y_{RESM} - RES) - K_{RESD}RES$$

$$\frac{dTKPD1}{dt} = F_{ProTKPD1}F_{THTKPD1}F_{IL2TK} - F_{TKPD1TKNPD1} - F_{TKPD1TEX} - F_{DTKPD1}$$

$$\frac{dTKNPD1}{dt} = F_{ProTKNPD1}F_{THTKNPD1}F_{IL2TK} + F_{TKPD1TKNPD1} - F_{DTKNPD1}$$

$$\frac{dTH}{dt} = F_{ProTH}F_{CANTH}F_{TREGTH} - F_{DTH}$$

$$\frac{dTREG}{dt} = F_{ProTREG}F_{CAFTREG} - F_{DTREG}$$

$$\frac{dTEX}{dt} = F_{ProTEX} + F_{TKPD1TEX} - F_{DTEX}$$

$$\frac{dFWT}{dt} = F_{ProFWT} + F_{CAFFWT} - F_{FWTCAF} - F_{DFWT}$$

$$\frac{dCAF}{dt} = F_{ProCAF}F_{M2CAF}F_{CANCAF}F_{OPNCAF} + F_{FWTCAF} - F_{CAFFWT} - F_{DCAF}$$

$$\frac{dMACM1}{dt} = F_{ProMACM1}F_{CANMACM1} + F_{M2M1} - F_{M1M2} - F_{DMACM1}$$

$$\frac{dMACM2}{dt} = F_{ProMACM2}F_{CAFMACM2} + F_{M1M2} - F_{M2M1} - F_{DMACM2}$$

$$\frac{dIL2}{dt} = F_{TKIL2} - F_{DIL2} + C_{IL2}$$

$$\frac{dLIF}{dt} = F_{CANLIF} + F_{CAFLIF} - C_{LIFKO}LIF$$

$$\frac{dIFNG}{dt} = F_{TKIFNG}F_{OPNIFNG} - F_{DIFNG}$$

$$\frac{dIL8}{dt} = F_{CANIL8} + F_{CAFIL8} + F_{M2IL8} - F_{DIL8} - C_{IL8KO}IL8$$

$$\frac{dLAC}{dt} = F_{CANLAC} + F_{MACLAC} - F_{DLAC} - C_{LIFKO}LAC$$

$$\frac{dIL10}{dt} = F_{TKIL10} - F_{DIL10}$$

$$\frac{dICAM1}{dt} = F_{TKICAM1} - F_{DICAM1}$$

$$\frac{dOPN}{dt} = (F_{CANOPN} + F_{CAFOPN})F_{IRF8OPN} - F_{DOPN} - C_{OPNKO}OPN$$

$$\frac{dIRF8}{dt} = F_{MACM1IRF8} - F_{DIRF8}$$

#### Abbreviations

Res = Resource concentration

**C_0 subtypes:**
CST = Killer T cell-exposed Tumor stem cells
CSNT = Non-Killer T cell-exposed Tumor stem cells

**C- subtypes:**
CNPDL1 = Tumor cells exposed to Killer T cells without PDL-1
CRNPDL1 = Tumor cells hidden from Killer T cells without PDL-1

**C+ subtypes:**
CPDL1 = Tumor cells exposed to Killer T cells with PDL-1
CRPDL1 = Tumor cells hidden from Killer T cells with PDL-1

**T cell subtypes:**
TKPD1 = Killer T cells with PD-1 (TK+)
TKNPD1 = Killer T cells without PD-1 (TK-)
TH = Helper T cells
TREG = Regulatory T cells
TEX = Exhausted T cells

**Fibroblast subtypes:**
FWT = Wild type fibroblasts (F_WT)
CAF = Invasive cancer associated fibroblasts

**Macrophage subtypes:**
MACM1 = M1 phase macrophage
MACM2 = M2 phase macrophage

**Molecular species:**
IL2 = Interleukin 2
LIF = Leukemia inhibitory factor
IFNG = Interferon gamma
IL8 = Interleukin 8
LAC = Lactate
IL10 = Interleukin-10
OPN = Osteopontin
IRF8 = Interferon regulatory factor 8

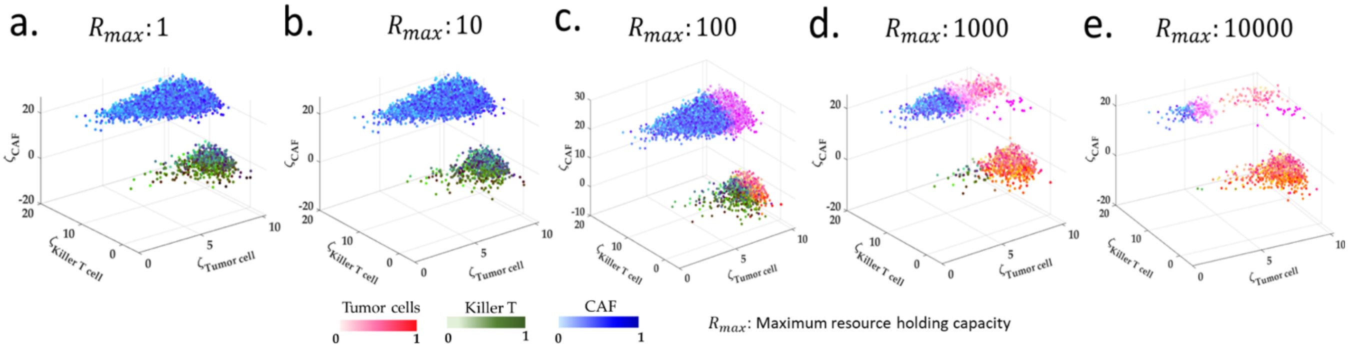


**Fig S2. Resource competition leads to different outcomes for different resource capacity. (a-b)** Under low-moderate resource availability, the parameter scan for the entire TME system reveals three possible TME composition Immune-dominated, fibro-desert, and desert. Interesting for a fixed resource supply, the parameter scan shows all the five TME subtypes. **(c-e)** The fibro-dominated and immune-desert subtypes emerge beyond a cut-off maximum resource supply rate along with the three mentioned subtypes. The high availability of resource ensures a constant amount of resource to all the different tumor cell types. Further, the elimination of PDL1- tumor cells by killer PD1+ T cells provide a competitive advantage to the PDL1+ tumor cells which, in turn enhances the T-cell exhaustion rate rendering a non-empty immune-desert region.

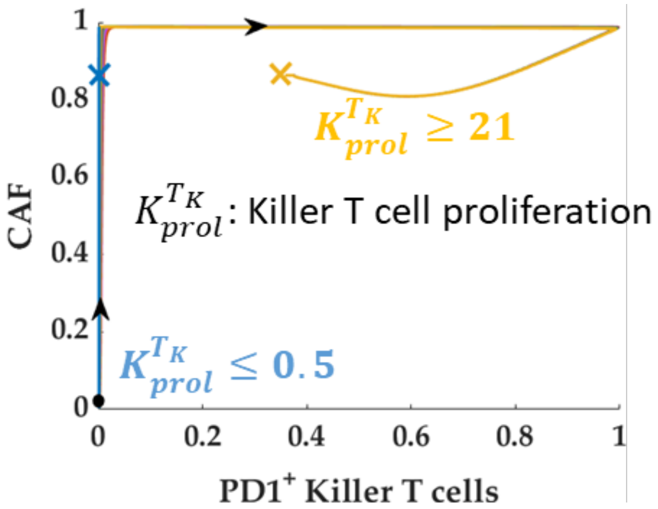


**Fig S3. Killer T cell-independent growth of CAF.** The proliferation rate governs the pre-ICI population of killer T cells. Below a critical proliferation rate the HNSCC TME model settles in an immune-desert region. Whereas, in both the scenarios (immune-desert and immune-rich), the CAF population remains unaltered indicating a relative independence from the T cell population.

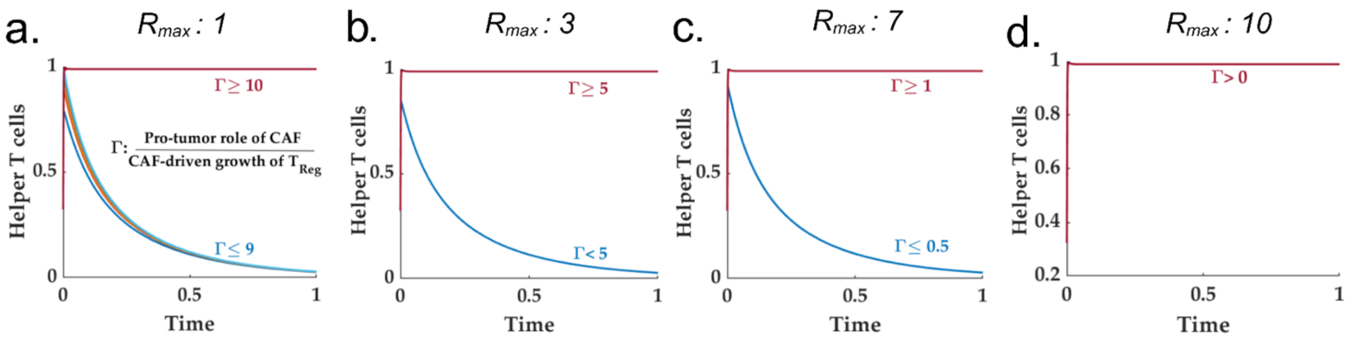


**Fig S4. Resource intake governs overall dependence of helper T cell on pro-tumor role of CAF. (a-d)** The pro-tumor role of CAF leads to significant pre-ICI, PDL1- tumor cell population. Therefore, beyond a threshold value of the pro-tumor role of CAF (compared to the CAF-driven growth of regulatory T cells), the final helper T cell population remains high. On the other hand, for moderate to low CAF-tumor interaction, the PDL1- tumor cells remain low due to the presence of cytotoxic killer T cells. Therefore, despite an initial increase, the helper T cells settle to a very low value (almost zero). Further, the threshold value of CAF-tumor interaction is dependent on the maximum resource intake in a competitive setting.

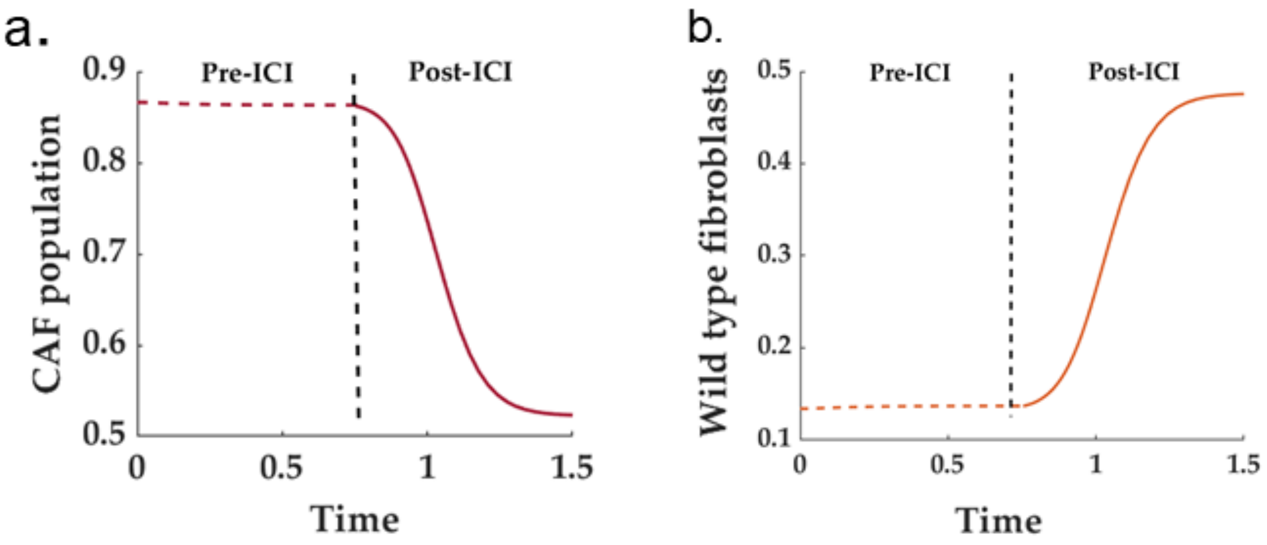


**Fig S5. ICI reduces CAF population and increases the wild type fibroblasts in immune rich scenario. (a)** The CAF population exhibits a steep increasing tendency owing to multiple paracrine interaction with the tumor cells and tumor associated macrophages. However, the ICI intervention in an immune rich scenario reduces the tumor cells. Further, the reduction in tumor cells-secreted LIF reduces the transition flux from wild type to cancer-associated fibroblasts. Therefore, overall CAF population undergoes a significant reduction during the ICI therapy. **(b)** The wild-type fibroblast population, due to significant reduction in the transition flux towards CAF, increases during an ICI-based therapy in immune rich scenario.

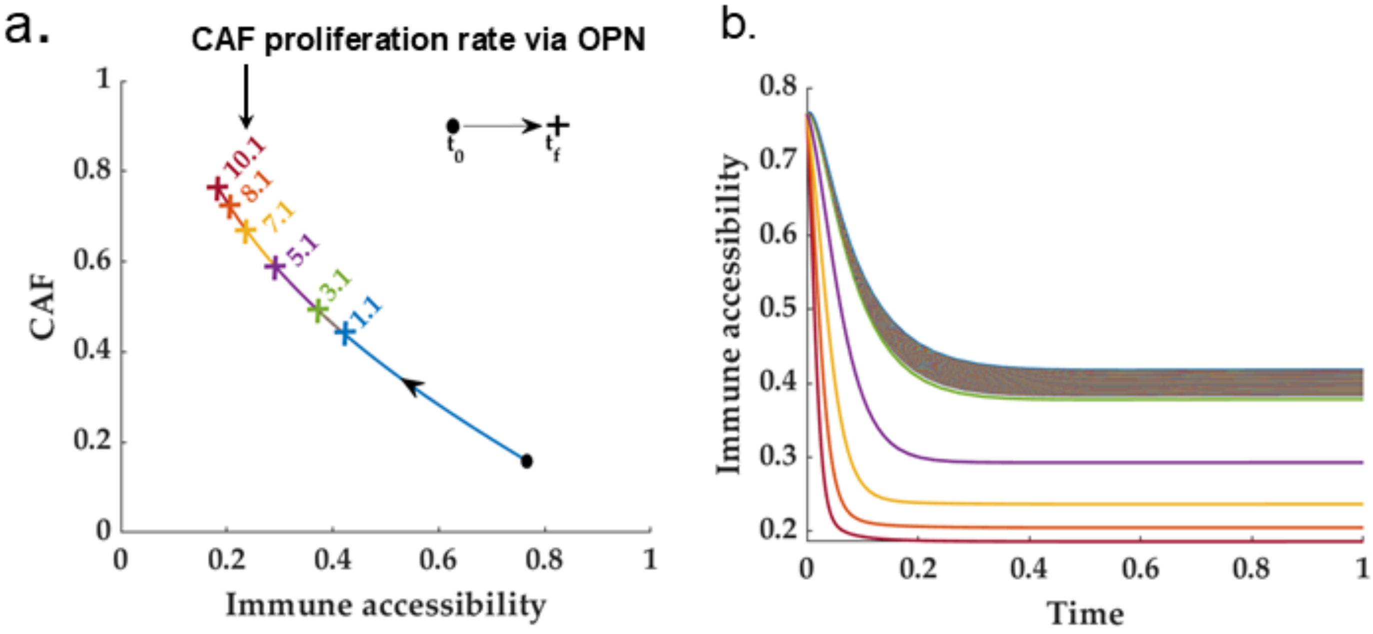


**Fig S6. The CAF-immune accessibility story. (a)** Demonstrates the phase-space between the immune accessibility and CAF for different CAF proliferation rate. **(b)** The time profile for immune accessibility shows the existence of a threshold time beyond which the immune accessibility deteriorates drastically. Further, this threshold time is dependent on the proliferation rate of CAF. This is due to the fact that a higher proliferation rate renders a faster CAF growth and due to the near-linear trajectory of CAF-immune accessibility trajectory, the immune accessibility adopts a faster time scale.

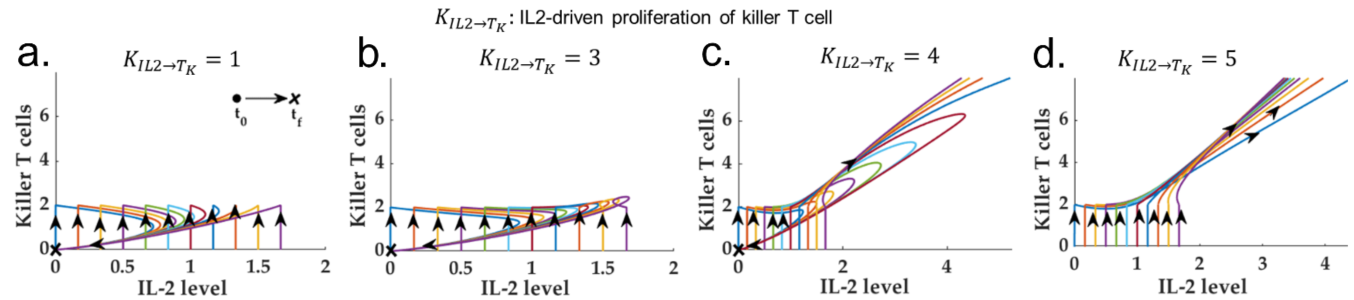


**Fig S7. IL2-Killer T cell story. (a-d)** Increasing levels of IL-2-induced killer T cell proliferation rate can drive the HNSCC immune-desert TME to an immune hot scenario. However, there exists a threshold IL-2-driven Killer T cell proliferation rate below which the immune-desert scenario can not be circumvented irrespective of the external IL-2 level.

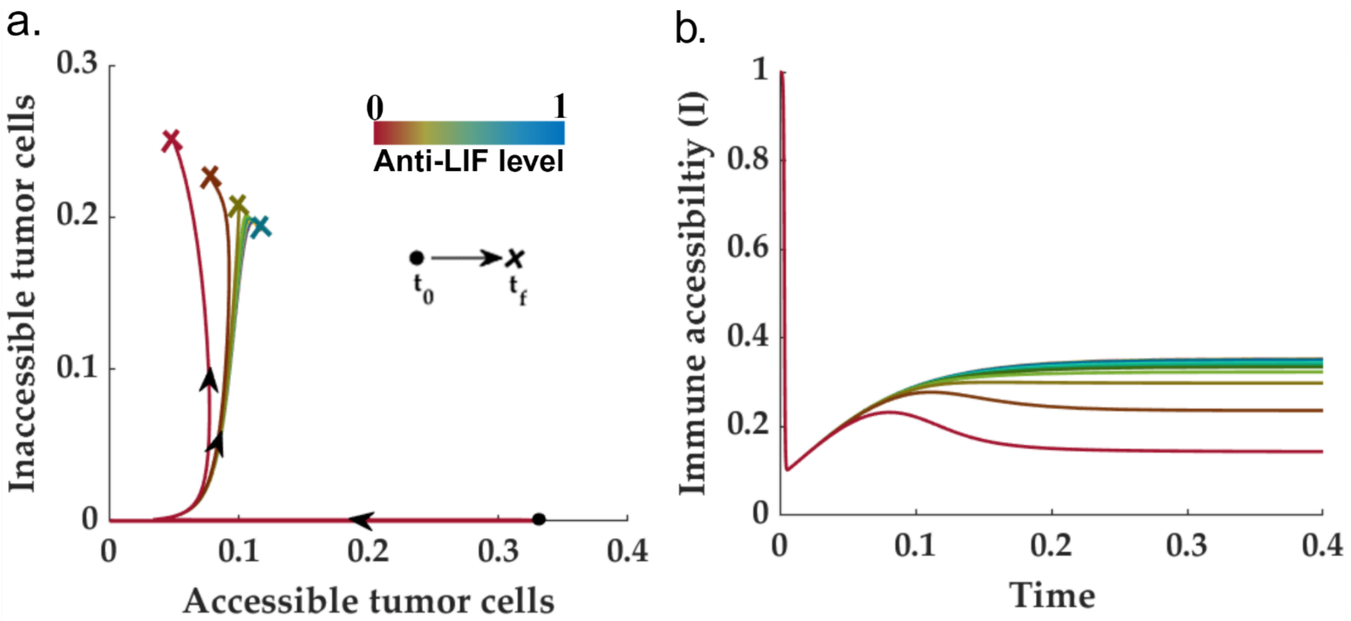


**Fig S8. LIF knockout reduces increases immune accessibility. (a)** Phase trajectory of different tumor cells vis-à-vis immune accessibility. The LIF knockout significantly reduces the pre-ICI inaccessible tumor cells. However, a complete (or near) complete elimination of inaccessible tumor cells is not possible with only LIF knockout. **(b)** Although a LIF knockout improves the immune accessibility it does not drive the TME system to an immune-dominated situation.

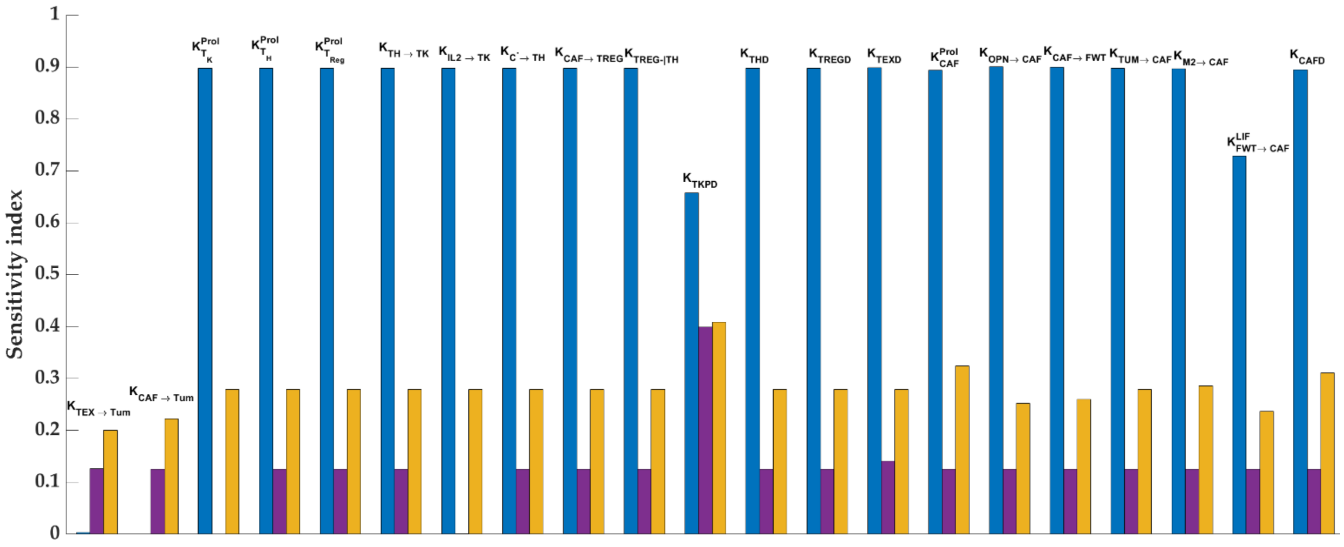

**Fig S9. Sensitivity analysis.** We chose all the parameters that exhibits an explicit bearing with the proliferation and death and conversion fluxes for Tumor cells (Blue), Killer T cells (Violet), and CAF (Yellow).

### Self-assessment of adherence to the Ten Simple Rules of Credible Practice in Modeling and Simulation in Healthcare

The current self-assessment of the manuscript titled, 'Tumor microenvironment governs the prognostic landscape of immunotherapy for head and neck squamous cell carcinoma: A computational model-guided analysis.' is in accordance with Erdemir et al. (2020). The rubric can be accessed at: https://www.imagwiki.nibib.nih.gov/content/10-simple-rules-conformance-rubric

Date of self-assessment: August 29, 2024

**Model files and documentation:** Provided in the supplementary text.

**Rule 1: Define context clearly:** Develop and document the subject, purpose, and intended use(s) of the model or simulation.

**Current Conformance Level:** Comprehensive

**Model Context:** Cell-state-specific mechanistic model of the tumor microenvironment (TME) for head and neck squamous cell carcinoma (HNSCC) in the presence (absence) of Immune checkpoint inhibitor treatment (ICI).

**Primary goal of the model/tool/database:** The primary goal of the modeling exercise was to leverage the TME-wide mechanistic models to explain (a) the existence of distinct compositional possibilities of the HNSCC TME and (b) how these compositional possibilities play a governing role in determining the response to ICI therapy. Additionally, the proposed model predicts the potential targets and biomarkers towards an improved ICI response.

**Biological Domain of the Model:** Cellular state

**Structures of the Model**: Tumor microenvironment

**Spatial Scales Included in the Model:** N/A (Assumes spatial homogeneity)

**Time Scales Included in the Model:** Week-Month

**<u>Rule 2: Use contextually appropriate data:</u>** Employ relevant and traceable information in the development or operation of a model or simulation.

**Current Conformance Level:** Adequate

| **Data for building the model** | **Published?** | **Private?** | **How is credibility checked?** | **Current Conformance Level** |
| --- | --- | --- | --- | --- |
| in vitro (primary cells cell, lines, etc.) | N/A | N/A | N/A | N/A |
| ex vivo (excised tissues) | N/A | N/A | N/A | N/A |
| in vivo pre-clinical (lower-level organism or small animal) | N/A | N/A | N/A | N/A |
| in vivo pre-clinical (large animal) | N/A | N/A | N/A | N/A |
| Human subjects/clinical | Yes | No | The source data is qualitative and the related clinical protocols have been published in peer-reviewed journals | Adequate |

| **Data for validating the model** | **Published?** | **Private?** | **How is credibility checked?** | **Current Conformance Level** |
| --- | --- | --- | --- | --- |
| in vitro (primary cells cell, lines, etc.) | N/A | N/A | N/A | N/A |
| ex vivo (excised tissues) | N/A | N/A | N/A | N/A |
| in vivo pre-clinical (lower-level organism or small animal) | N/A | N/A | N/A | N/A |
| in vivo pre-clinical (large animal) | N/A | N/A | N/A | N/A |
| Human subjects/clinical | Yes | No | The source data is qualitative and the clinical protocols have been published in peer-reviewed journals | Adequate |

**Rule 3: Evaluate within context:** Perform verification, validation, uncertainty quantification, and sensitivity analysis of the model or simulation with respect to the reality of interest and intended use(s) of the model or simulation.

**Current Conformance Level:** Extensive

| | **Who Does It?** | **When does it happen?** | **How is it done?** | **Current Conformance Level** |
|---|---|---|---|---|
| **Verification** | Developer | During development | Comparison of model output with the experimental and clinical observations | Extensive |
| **Validation** | Lab Member | During development | model was used to reproduce simulations and figures | Extensive |
| **Uncertainty Quantification** | User performs uncertainty quantification | Can be performed every time the model is run for a new scenario | User discretion | Adequate |
| **Sensitivity Analysis** | User performs sensitivity analysis on influential parameters. | Can be performed after every new simulation | User discretion | Adequate |

**<u>Rule 4: List limitations explicitly:</u>** Provide restrictions, constraints, or qualifications for or on the use of the model or simulation for consideration by the users or customers of a model or simulation.

**Current Conformance Level:** Comprehensive

| **Disclaimer statement (explain key limitations)** | **Who needs to know about this disclaimer?** | **How is this disclaimer shared with that audience?** | **Current Conformance Level** |
|---|---|---|---|
| Models are limited by the spatial homogeneity approximation | Users | Stated in the main text | Comprehensive |
| Model does not capture metastasis and the associated transition and other necessary interactions | Users | Stated in the main text | Comprehensive |
| The conclusion drawn from this model are conditioned on the particular modeling rules specified in the main text | Users | Stated in the main text | Comprehensive |

**<u>Rule 5: Use version control:</u>** Implement a system to trace the time history of modeling and simulation activities including delineation of each contributors' efforts.

**Current Conformance Level:** Extensive

| | **Naming Conventions?** | **Repository?** | **Code Review?** |
|---|---|---|---|
| **individual modeler** | N/A | Github | Yes |
| **within the lab** | Yes | Yes | Yes |
| **collaborators** | N/A | Github | Yes |

**<u>Rule 6: Document appropriately:</u>** Maintain up-to-date informative records of all modeling and simulation activities, including simulation code, model mark-up, scope and intended use of modeling and simulation activities, as well as users' and developers' guides.

**Current Conformance Level:** Extensive

| | **Current Conformance Level** |
|---|---|
| **Code Commented?** | Extensive: Commented the codes in every important line of the code for better interpretability. |
| **Scope and intended use described?** | Extensive: Described in the beginning of the each section of the code and in the main text. |
| **User's Guide** | Extensive: described in the main text and supplemental files |
| **Developer's Guide?** | Adequate: Described in the main text and the supplementary files. |

**<u>Rule 7: Disseminate broadly:</u>** Share all components of modeling and simulation activities, including simulation software, models, simulation scenarios and results.

**Current Conformance Level:** Extensive

| **Target Audience(s):** | **"Inner Circle"** | **Scientific Community** |
|---|---|---|
| **Simulations** | Shared with the lab members for replication | The co-authors of this work presented this in the form of posters in FOSBE2024, VPH2024 |
| **Models** | Shared with the lab members for replication | The co-authors of this work presented this in the form of posters in FOSBE2024, VPH2024 |
| **Software** | MATLAB is well-known software in the lab of the first author. | MATLAB is well-known software in the mathematical modeling community. |
| **Results** | Presented at lab members | MATLAB is well-known software in the lab of the first author. |
| **Implication of Results** | Presented at lab members | MATLAB is well-known software in the lab of the first author. |

**<u>Rule 8: Get independent reviews</u>:** Have the modeling and simulation activity reviewed by nonpartisan third-party users and developers.

**Current Conformance Level:** Extensive

| **Reviewer(s) name and affiliation** | **Alexandra Manchel (Thomas Jefferson University)** |
|---|---|
| When was the review performed? | August 15, 2024 |
| How was review performed and outcomes of the review? | A member of the research group, not involved in the present study and does not conduct research in tumor microenvironment modeling, performed the review. Model scripts were cross-checked for consistency. Simulation results and figures were replicated using the files provided on GitHub. |

| **Reviewer(s) name and affiliation** | **Dr. Prem Jagadeesan (Purdue university)** |
|---|---|
| When was the review performed? | August 29, 2024 |
| How was review performed and outcomes of the review? | A member of the research group, not involved in the present study and does not conduct research in tumor microenvironment modeling, independently produced the Figure 3 of the manuscript from the model equations, parameters, and initial conditions provided on GitHub. |

**Rule 9: Test competing implementations**: Use contrasting modeling and simulation implementation strategies to check the conclusions of different strategies against each other.

**Current Conformance Level:** Adequate

| | **Yes or No (briefly summarize)** |
|---|---|
| **Were competing implementations tested?** | Competing implementations were conceptualized by all the authors and tested by the first author of this manuscript. |
| **Did this lead to model refinement or improvement?** | Yes, the completing implementations led to modifications and refinements of the model. |

**Rule 10: Conform to standards:** Adopt and promote generally applicable and discipline specific operating procedures, guidelines, and regulations accepted as best practices.

**Current Conformance Level:** Adequate

| | **Yes or No (briefly summarize)** |
|---|---|
| **Are there operating procedures, guidelines, or standards for this type of multiscale modeling?** | Yes, the existing modeling approaches for tumor microenvironment employs continuous-time mathematical models which can vary between deterministic and stochastic settings. |
| **How do your modeling efforts conform?** | The proposed model adopts a continuous time modeling framework in a deterministic setting. Therefore, the modeling framework conforms to the existing conventions in the modeling literature. |
